# Reprogrammed Serine Integrases Enable Precise Integration of Synthetic DNA

**DOI:** 10.1101/2024.05.09.593242

**Authors:** Friedrich Fauser, Sebastian Arangundy-Franklin, Jessica E Davis, Lifeng Liu, Nicola J Schmidt, Luis Rodriguez, Danny F Xia, Nga Nguyen, Nicholas A Scarlott, Rakshaa Mureli, Irene Tan, Yuanyue Zhou, Lynn N Truong, Sarah J Hinkley, Bhakti N Kadam, Stephen Lam, Bryan Bourgeois, Emily Tait, Mohammad Qasim, Vishvesha Vaidya, Adeline Chen, Andrew Nguyen, Yuri R. Bendaña, David A. Shivak, Patrick Li, Andreas Reik, David E Paschon, Gregory D Davis, Jeffrey C Miller

## Abstract

Despite recent progress in the ability to manipulate the genomes of eukaryotic cells^1–3^, there is still no effective and practical method to precisely integrate large synthetic DNA constructs into desired chromosomal sites using a programmable integrase. Serine integrases can perform the necessary molecular steps^4^, but only if their natural target site is first installed into the recipient genome by other methods. A more elegant approach would be to directly reprogram the serine integrase itself to target a desired genomic site that is different from the natural recognition site of the integrase^5^. Here, we describe the development of a Modular Integrase (MINT) platform, a versatile protein-guided genome editing tool that can facilitate site-directed targeted integration (TI) of synthetic DNA into chromosomal sites. Through a combination of structural modeling, directed evolution, and screening in human cells we have reprogrammed the specificity of the serine integrase Bxb1. We demonstrate the therapeutic potential of the MINT platform by retargeting Bxb1 to the human AAVS1 and TRAC loci where wild-type Bxb1 has no detectable activity. By combining the MINT platform with known activity-increasing Bxb1 mutants, we achieved 14% TI at the AAVS1 locus, and by additionally fusing zinc finger DNA binding domains to engineered Bxb1 variants, we achieved 35% TI at the TRAC locus in human K562 cells. To further demonstrate clinical potential, we achieved stable 15% TI of a functional donor construct in human T cells.

## Introduction

The ability to integrate synthetic DNA constructs into desired chromosomal locations in eukaryotic genomes would have broad implications for the development of genomic medicines, agriculture, synthetic biology, and basic research. Initially, there was excitement that homology-directed-repair (HDR) stimulated by engineered nucleases would be able to accomplish this goal, but numerous limitations were encountered when this was assessed, such as low efficiency caused by competing DNA repair pathways, undesired chromosomal rearrangements, and break-induced DNA damage response^6–9^. Furthermore, despite the significant progress in programmable DNA binding using zinc fingers (ZFs)^10–12^, transcription activator-like effectors (TALEs)^13,14^, and RNA-guided proteins such as Cas9^15^, simply tethering such engineered DNA binding domains to the catalytic domains of recombinases or transposons has not yielded reagents capable of driving high levels of integration in human cells^16–23^. We initially explored such systems and observed high levels of indels (Extended Data Fig. 1a) which may partially explain the difficulties others have encountered.

Greater success has been achieved by inserting ZFs into the tyrosine recombinase Cre to perform inversions^24^, however fusions to other types of integrases have not proved nearly as successful^25–29^. There has been some recent progress using CRISPR-associated transposases^30–33^ but delivering such complex systems to therapeutically relevant cells has yet to be demonstrated and other nucleic acid-guided systems have target site size limitations^34^ that will make specifically targeting therapeutically relevant loci difficult.

Large serine recombinases (LSRs - also known as Lare Serine Integrases, LSIs) have long been proposed as the ideal tool for genome engineering^35,36^ due to their unique properties including irreversible integration and no inherent limit to the size of the DNA constructs they can integrate (**Figure 1a**). In their natural context, an integrase dimer binds to a phage attP site, while another dimer binds an attB site in the bacterial host’s genome. The assembled and fully active tetramer then forms a covalent bond to the DNA via its active-site serine residues producing a temporary two base-pair overhang in the center of each binding site which is referred to as the central dinucleotide or CDN. If the sequence of the CDN in the attP site matches the sequence of the CDN in the attB site then subunit rotation facilitates strand exchange and ligation, without leaving free DNA ends or nicks, thus obviating the need to engage the host cell’s DNA repair mechanisms. Importantly, in the absence of a phage-encoded reversibility factor, integration proceeds unidirectionally and is irreversible. Nevertheless, the use of LSRs as genome editing reagents has been hampered by the difficulty in their engineering and deployment towards non-cognate targets.

**Figure 1.**
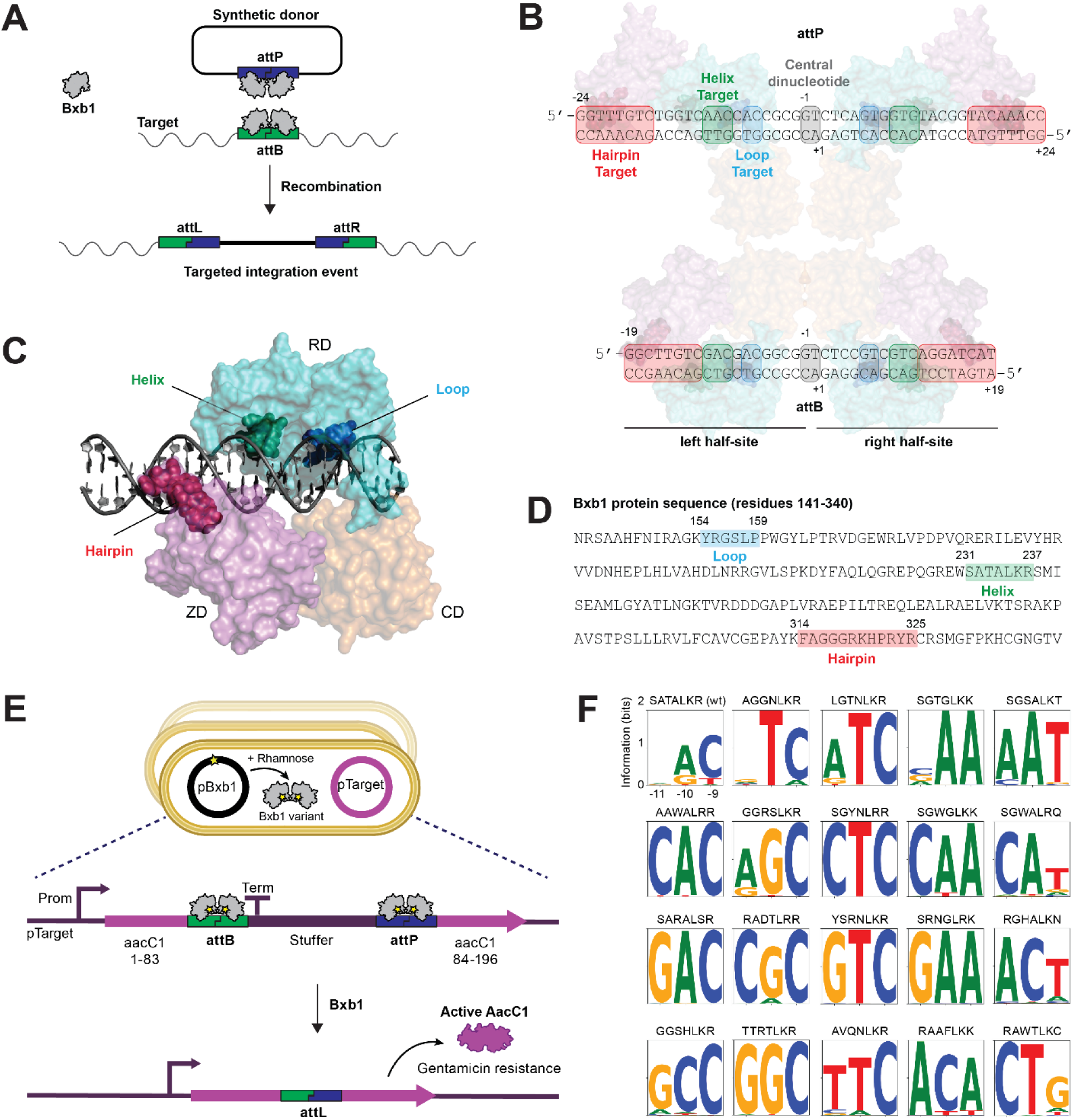
Bxb1-DNA interaction mapping and DNA binding domain engineering using directed evolution. **a.** Schematic of Bxb1-induced TI of a synthetic circular donor into a genomic target. Attachment sites are bound by Bxb1 dimers and non-reversible recombination is facilitated by a Bxb1 tetramer. **b.** The sequence of the natural Bxb1 attP and attB attachment sites with the portions recognized by the hairpin, helix, and loop regions of Bxb1 highlighted. The central dinucleotide of the attachment sites is not directly recognized by Bxb1 but needs to match between the attP and attB site for integration to occur. **c.** Structural model of a Bxb1 monomer bound to an attB attachment half-site. **d.** Amino acid sequence of wild-type Bxb1 with residues investigated in this study highlighted. Bxb1-DNA interaction mapping identified three specificity-determining regions of Bxb1 that can be reprogrammed: the loop region (residues 154-159; shown in blue), the helix region (residues 231-237; shown in green) which are both in the RD domain of Bxb1, and the hairpin region (residues 314-325; shown in red) within the ZD domain of Bxb1. **e.** Schematic of our two-plasmid directed evolution system. The expression plasmid pBxb1 encodes an integrase gene, while the reporter plasmid pTarget contains an antibiotic marker disrupted by a stuffer sequence that is flanked by modified attB and attP sequences. Recombination of these sites by an active integrase excises the stuffer sequence and restores the open reading frame of the antibiotic resistance marker. The selection system using gentamicin resistance is shown. The T in the stuffer sequence indicates the position of a transcriptional terminator sequence. This system enables either screening of libraries of integrase variants against a single DNA target or screening of a library of DNA targets against a single integrase. **f.** Example DNA specificity plots of selected helix variants. Data in panel d is the mean of four biological replicates. Plotted data are provided in the Source Data file.

The best results to date have been achieved with systems that use a phage-derived serine integrase such as Bxb1 combined with prime editing to simultaneously introduce the natural attB or natural attP target site into mammalian cells^37–39^. However, the requirement to first use prime editing to integrate the target site makes the process more complex for therapeutic development and manufacturing, requiring the simultaneous delivery of two independent genome editing systems to the cell. A more straightforward approach would involve directly reprogramming the target preference of the serine integrase itself while retaining the desirable properties of natural LSRs.

Despite the utility of reprogramming an LSR such as Bxb1^40^, there are several technical challenges to overcome. While TALEs and CRISPR systems can easily be targeted to desired DNA sequences using simple targeting rules, no one has been able to divine such a “simple code” that governs Bxb1 retargeting. Directed evolution systems have been used to engineer proteins such as ZFs that lack a simple DNA targeting code, but the modularity and compactness of the ZF repeat allows the engineering of one or two ZF repeats that can then be combined into a longer array^41,42^. Directed evolution can also be used on the entire coding sequence of non-modular proteins such as Cre, but this required over a hundred sequential rounds of directed evolution to achieve good performance at a desired target site^43,44^. A further challenge for reprogramming Bxb1 is that no experimentally determined structure of Bxb1 bound to its target site is currently available.

In this study, we used a combination of structural modeling and experimental characterization to map critical protein-DNA interactions between Bxb1 and its target site. The only known structures of LSRs bound to target DNA suggest that the zinc ribbon domain (ZD) and the recombinase domain (RD) are separated by a flexible polypeptide linker and recognize their portions of the attB and attP targets in a modular fashion^45^. On the assumption that Bxb1 recognizes its target site in a similar way, we developed a strategy for LSR engineering that uses directed evolution to reprogram the key specificity-determining amino acid residues of the RD and ZD domains in parallel and then combine the successful variants into fully reprogrammed LSR variants which we refer to as MINT constructs. We use this strategy to successfully target multiple pseudo-sites in the human genome where wild-type Bxb1 has detectable activity as well as intron 1 of the therapeutically relevant human AAVS1 and TRAC loci where wild-type Bxb1 has no detectable activity. We then add previously described mutations that increase the activity of wild-type Bxb1 to increase integration activity of our MINT constructs that target both AAVS1 and TRAC. Finally, we fuse engineered ZF arrays to our engineered LSRs that target the human TRAC locus to achieve 35% TI in human K562 cells and stably integrate a functional GFP-expressing donor with a frequency of 15% in primary human T cells.

### Bxb1-DNA interaction mapping

To compensate for the lack of a known structure of Bxb1 bound to its DNA target site, we performed an initial experiment designed to map interactions between residues in the Bxb1 RD and ZD domains and key regions of its natural DNA target site. Based on a sequence alignment of Bxb1 and the regions of the LI integrase known to interact with its DNA target^45,46^ (Extended Data Fig. 2), we identified 89 residues in Bxb1 that are likely to interact with DNA. We performed scanning mutagenesis on these 89 residues and co-delivered the resulting Bxb1 variants along with plasmids bearing variant attB and attP sites and measured recombination in human K562 cells. This experiment identified two clusters of residues which, when mutated, could alter specificity at portions of the DNA target site. A first cluster of residues at positions 231, 233, and 237 of Bxb1 was able to alter specificity at positions −10 and −9 of the DNA target site. A second cluster of residues at positions 314, 315, 316, 318, 323, and 325 of Bxb1 was able to alter specificity at positions −18 through −13 of the DNA target site. The residue at position 158 of Bxb1 was also able to alter specificity at positions −7 and −6 of the DNA target site. Saturation mutagenesis of these residues was able to achieve further specificity shifts. Mapping the Bxb1 mutations that shifted targeting specificity onto a RoseTTAFold^47^ structural model of Bxb1 bound to DNA indicated two regions in the RD domain and one region in the ZD domain where focused protein engineering should be able to alter the DNA targeting specificity. We refer to these regions as the helix (positions 231-237), hairpin (positions 314-325), and loop (positions 154-159) respectively based on the structure of each region in our Bxb1 structural model (**Figure 1b-d**).

### Evolving Bxb1-DNA interaction domains

Next, we developed a directed evolution system to engineer Bxb1 target site specificity. This system is based on a previously described method^48^ whereby a “stuffer” sequence flanked by artificial attB and attP target sites is placed within the coding sequence of an antibiotic resistance gene such that recombination by a Bxb1 variant able to target the attB and attP sites restores the open reading frame and allows the bacterial cell to survive an antibiotic challenge (**Figure 1e**; Extended Data Fig. 3a). In our system we placed the Bxb1 variant library and the DNA target sites on separate plasmids to enable more rapid testing of the same library of Bxb1 variants against different DNA target sites and we utilized a different antibiotic resistance gene to improve selection performance. A related system was used to assess the DNA targeting specificity of a selected Bxb1 variant by testing a library of different target sites against a single Bxb1 variant (Extended Data Fig. 3e).

Our structural analysis of the LI integrase indicated that residues in the structure corresponding to residues 231-237 in Bxb1 form an alpha helix that docks with its target DNA in a manner reminiscent of the interaction between a zinc finger (ZF) and its target trinucleotide^49^. Thus, we also adopted the same residue randomization scheme used with ZFs^42^ and kept residue 235 fixed as a leucine since this seems analogous to the leucine that is often at +4 of a ZF recognition helix. This resulted in a library of Bxb1 helix variants with residues 231-234, and 236-237 fully randomized (Extended Data Fig. 3a,b). Since ZFs can target 3 bp DNA sequences we hypothesized that the Bxb1 helix might be able to specify the DNA bases at positions −11, −10, and −9 of the target site. Thus, we performed 64 separate selections using our Bxb1 helix variant library against all 64 possible DNA triplets at positions −11, −10, −9 (and mirrored at positions +9, +10, +11) of the attB and attP target sites. Many selected helices demonstrated dramatic changes in target specificity at these positions in the DNA target site. A comparison of the target preference for the wild-type Bxb1 helix SATALKR and 19 selected helix sequences is shown in **Figure 1f** and a ranked list of 366 selected helices organized by their preferred DNA triplet can be found in Supplementary Tables 1,2.

Next, we performed selections using a library of Bxb1 variants where positions 154-159 (the loop region) were randomized (Extended Data Fig. 3a,c). We performed 16 separate selections using this library against all possible DNA dimer sequences at positions −7 and −6 and found that, surprisingly, a single residue change of S157G could shift the sequence preferences of wild-type Bxb1 towards nearly every single dinucleotide at positions −7 and −6 except for CG (Extended Data Fig. 3g) and we note that the same mutation was also identified by a recent effort to increase the activity of Bxb1^39^. Other selected loop variants were able to show improved target specificity relative to the wild-type loop, but the sequences that can be targeted specifically in the loop variants we have characterized tend to be limited to sequences with A or T at position −7 and C or T at position −6.

Finally, we used our directed evolution system to select Bxb1 variants with mutations at the hairpin region (residues 314-325). Since a fully randomized hairpin region would not be efficiently covered by a typical library generated in E. coli, we fully randomized residues 314, 316, 318, 321, 323, and 325 that partially face the DNA major groove in our model, and partially randomized residue 322 to correspond to residues conserved in many large serine recombinases (Extended Data Fig. 3a,d). Correspondingly, the DNA target sequence of the hairpin encompasses a sequence space too large to easily perform a selection against each possible target site variant so we performed selections against specific target sites where positions −19 to −12 matched potential Bxb1 target sites of interest in the human genome. Our selections returned hairpin motifs with wide sequence divergence (Extended Data Fig. 3f) and obvious target preference differences in comparison to the wild-type hairpin.

### Identifying Bxb1 pseudo-sites in the human genome

As an initial test of engineered Bxb1 variants in human cells, we decided to target Bxb1 pseudo-sites in the human genome where wild-type Bxb1 already has detectable activity so that even modest improvements could be measured. At the time, no human Bxb1 pseudo-sites were known so we attempted to identify such sites in human K562 cells by performing both a computational search based on published Bxb1 specificity data^36^ and a genome-wide experimental approach that mapped integrated donor constructs using a method similar to anchored multiplex PCR^50,51^ (Extended Data Fig. 4a). In order to comprehensively identify pseudo-sites, we ignored the CDN sequence in the computational approach and we used a mixture of donors with all possible 16 CDNs for the experimental approach.

Combining the results of both approaches, we were able to identify 23 sites with at least 0.1% integration in human K562 cells. We chose five sites with between 39% and 61% homology to the natural Bxb1 attB site where wild-type Bxb1 achieves between ∼0.20% and ∼2.45% integration as test targets for our engineered Bxb1 variants (Extended Data Fig. 4b, Supplementary Tables 3,4). Notably, both computational and experimental approaches identified only active attB pseudo-sites, and no attP pseudo-sites, mirroring the direction of natural Bxb1-directed integration into its host genome. This is consistent with our finding that “landing pad” cell lines with attB sites pre-integrated into the genome support higher levels of integration than landing pad cells lines with attP sites pre-integrated (Extended Data Fig. 1b).

### Increasing Bxb1 activity at human pseudo-sites

Having identified suitable Bxb1 pseudo-sites in the human genome, we decided to test if selected Bxb1 helix variants could improve activity at an intergenic pseudo-site on chromosome 3 (Extended Data Fig. 5a). This pseudo-site has one half-site with a T at position −10 that is expected to be recognized poorly by the wild-type Bxb1 helix (**Figure 1f**). To improve activity, we performed a helix selection against the relevant portion of this half-site and the selection yielded multiple families of helix sequences. We tested a total of 58 selected helices from these different sequence families for their ability to carry out site-specific integration in human K562 cells. Data for eight helices that represent the most active variants from each sequence family with activities ranging from 8% to over 30% are shown in Extended Data Fig. 5a. Note that the selected helix AGGNLKR demonstrates a dramatic shift in DNA preference towards a T at position −10 (**Figure 1f**). Next, we wanted to investigate whether helix and hairpin variants, located in separate domains of Bxb1, can be engineered independently and then combined to further engineer Bxb1 towards this target site. We performed a selection using our Bxb1 variant hairpin library against the relevant portion of the same half-site of the chromosome 3 target and identified the hairpin peptide LARGRRKWARYR that can further improve performance when combined with selected helix variants (Extended Data Fig. 5b). The combination of the AGGNLKR helix and LARGRRKWARYR hairpin yielded a Bxb1 variant with a substantial shift in genome-wide DNA target site specificity relative to wild-type Bxb1. For this variant, the intended pseudo-site in chromosome 3 has the most integrations within the entire human genome (Extended Data Fig. 5b) as assayed by an improved genome-wide specificity assay (Extended Data Fig. 6). Lastly, we confirmed that our Bxb1 variants are also active in other cell types, such as human HepG2 cells (Extended Data Fig. 5c).

We hypothesized that engineering separate variants for the left and right half-site of a pseudo-site could yield additional improvements in performance. But we were concerned that simply testing variants at the full endogenous pseudo-site would make it difficult to determine how recognition of each half site was being effected by a given Bxb1 variant so we established a system that utilizes synthetic DNA target sites on plasmids where recombination at an inverted repeat of a given left half-site could be monitored separately from recombination at an inverted repeat of a given right half-site. This system also allows us to test the activity of Bxb1 helix, loop, or hairpin variants against hybrid half-sites where the portion of the half-site recognized by either the RD or ZD domain can be replaced with the relevant portion of the wild-type Bxb1 attB target site (we refer to these as “quarter-sites”). Helix and/or loop variants that perform well at an RD quarter-site can then be combined with hairpin variants that perform well at the corresponding ZD quarter-site in order to recognize the desired half-site (**Figure 2a**). We used this approach with two endogenous pseudo-sites identified by the computational genome scan and two endogenous pseudo-sites identified by the experimental genome-wide screen. This system allowed us to rapidly identify evolved helix and hairpin variants that, when combined, are active against their desired half-sites (**Figure 2b,c**; Extended Data Fig. 5d-f), demonstrating the modularity of our MINT platform and we refer to such combinations of Bxb1 variants in the RD and ZD domains as MINT constructs. We were then able to achieve substantial improvements in activity relative to wild-type Bxb1 at 6 half-sites from these four human pseudo-sites (Extended Data Fig. 5g).

**Figure 2.**
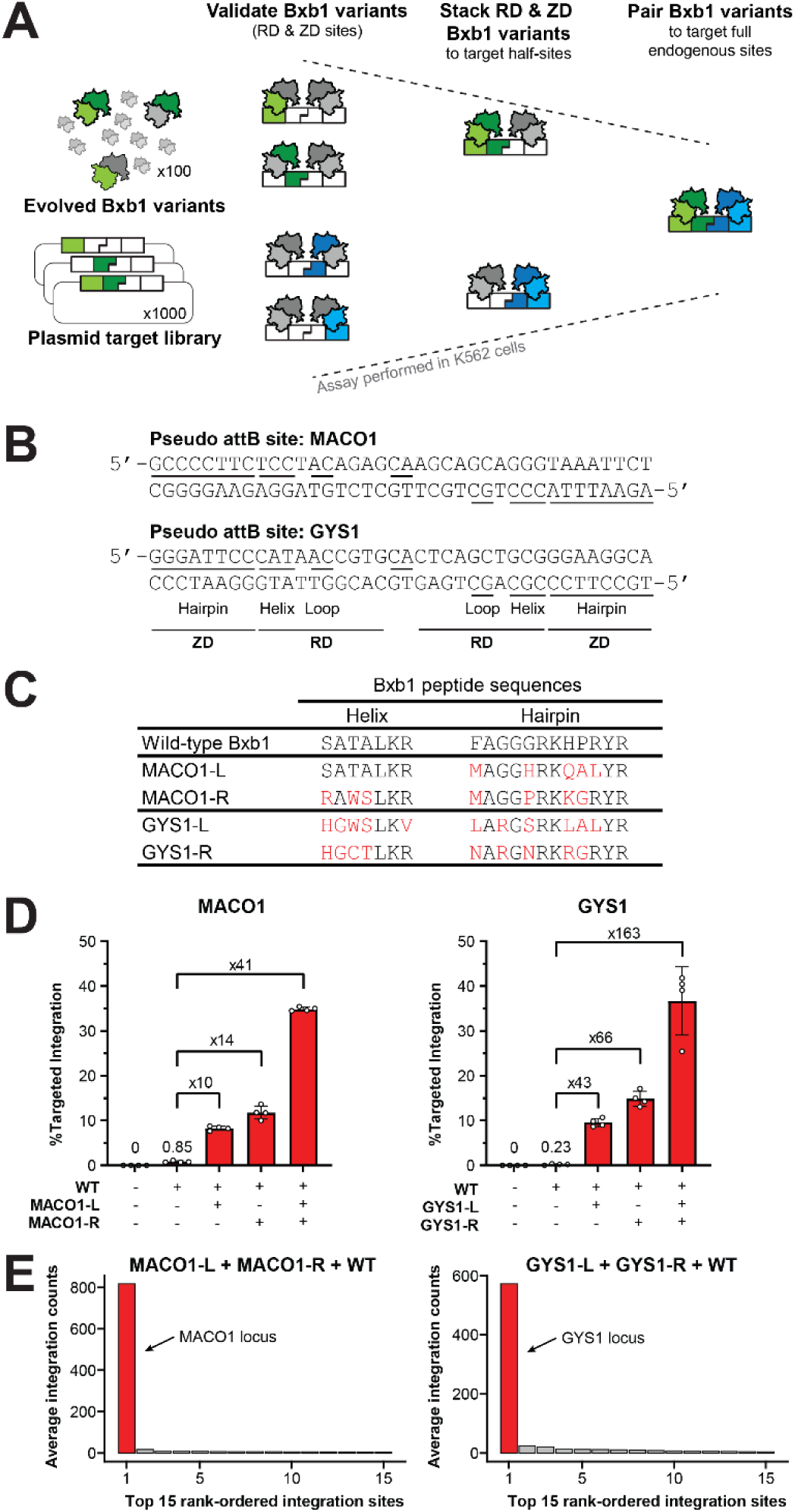
Performance of engineered Bxb1 variants at pseudo-sites in the human genome. **a.** Schematic of plasmid-based system for testing evolved Bxb1 variants against artificial DNA targets. Our LSR engineering strategy divides each endogenous target site into four “quarter-sites” where the left and right half-sites are each further divided into the portion recognized by the RD domain and the portion recognized by the ZD domain of Bxb1. Individual selected Bxb1 variants are screened against a library of plasmid targets that includes their intended quarter-sites. Successful RD and ZD variants are then combined and tested against the same library of plasmid targets that also includes the relevant half-sites. A single plasmid target library can contain quarter-site and half-site targets for numerous full endogenous target sites. Left and right site candidates derived from this assay can then be tested as pairs against the endogenous target site. See Extended Data Fig. 7b,c for additional details. **b.** Sequence of two Bxb1 pseudo-sites in the human genome. Both sites were identified experimentally using wild-type Bxb1. **c.** Bxb1 peptide sequences of evolved Bxb1 variants that showed improved performance against their corresponding half-sites. **d.** Results from a PCR-based NGS assay demonstrating improved performance of evolved Bxb1 variants against their chromosomal endogenous targets in K562 cells. The presence of a wild-type Bxb1 expression construct is necessary to bind the wild-type attP sequence on the donor plasmid. Data are presented as the mean ± s.d. from four biological replicates. **e.** Results from an unbiased genome-wide specificity assay demonstrating a strong on-target preference of our engineered Bxb1 variants. The top 15 candidate integration sites are shown (out of total 112 sites for MACO1 constructs and 103 for GYS1 constructs) with the entire datasets provided in Supplementary Tables 5 and 6 and corresponding control datasets in Supplementary Tables 7 and 8. Average integration counts were determined using the average deduplicated read numbers across 2 replicates and 2 samples each using primers that amplify the attL and attR plasmid-genome junctions. Plotted data are provided in the Source Data file.

Four of these half-sites comprise full pseudo-sites in the *MACO1* and *GYS1* genes and activity at these sites could be increased further by pairing the two MINT constructs that worked best at the corresponding left and right half-sites, achieving ∼35% TI at both loci (**Figure 2d**). For *MACO1* we observed 0.85% TI with wild-type Bxb1 and achieved a 41-fold increase in activity with a combination of our *MACO1* MINT constructs, while we detected 0.23% TI with wild-type Bxb1 at *GYS1* and achieved a 163-fold increase with a combination of our *GYS1* MINT constructs. We then investigated the genome-wide specificity of these two sets of MINT constructs (**Figure 2e**; Supplementary Tables 5-8). The top candidate integration site for both variants was the intended locus, with the signal for all other sites at least 20-fold lower. We did observe some integration at the *MACO1* pseudo-site with the *GYS1* MINT constructs, but this appeared as #13 in the candidate integration site list with 100-fold lower signal than the intended *GYS1* target. Conversely, integration at the *GYS1* target was not observed for the *MACO1* MINT constructs. Thus, we were able to retarget Bxb1 to previously weakly-active pseudo-sites with active and specific novel Bxb1 variants, indicating we had accomplished proper retargeting and did not simply relax the specificity of the parental constructs.

### Targeting clinically-relevant sites in the human genome

Encouraged by our success targeting chromosomal pseudo-sites with our MINT constructs, we proceeded to the more difficult challenge of targeting clinically relevant regions of the human genome, such as the well-established safe harbor AAVS1 locus^52^ and the T-cell receptor α constant (TRAC) locus (**Figure 3a**), in which wild-type Bxb1 has no detectable activity. We first screened Bxb1 variants against a library of potential target sites within intron 1 of these regions using our mammalian system (Extended Data Fig. 7, Supplementary Tables 9-13). We then performed custom hairpin selections for promising sites derived from this initial screen. This resulted in two pairs of MINT constructs (**Figure 3b**) that can recognize the AAVS1 and TRAC loci respectively and that each achieved up to ∼1% TI (**Figure 3c,d**; Extended Data Fig. 8,9). We utilized DNA donors with wild-type attP target sites with CDNs matched to the intended genomic target site and co-delivered wild-type Bxb1 to recognize the attP site in the donor. Notably, we did not observe any integration activity above background levels when we omitted the Bxb1 variant targeted to either the left or right half-site, indicating full retargeting of our MINT constructs to TRAC and AAVS1.

**Figure 3.**
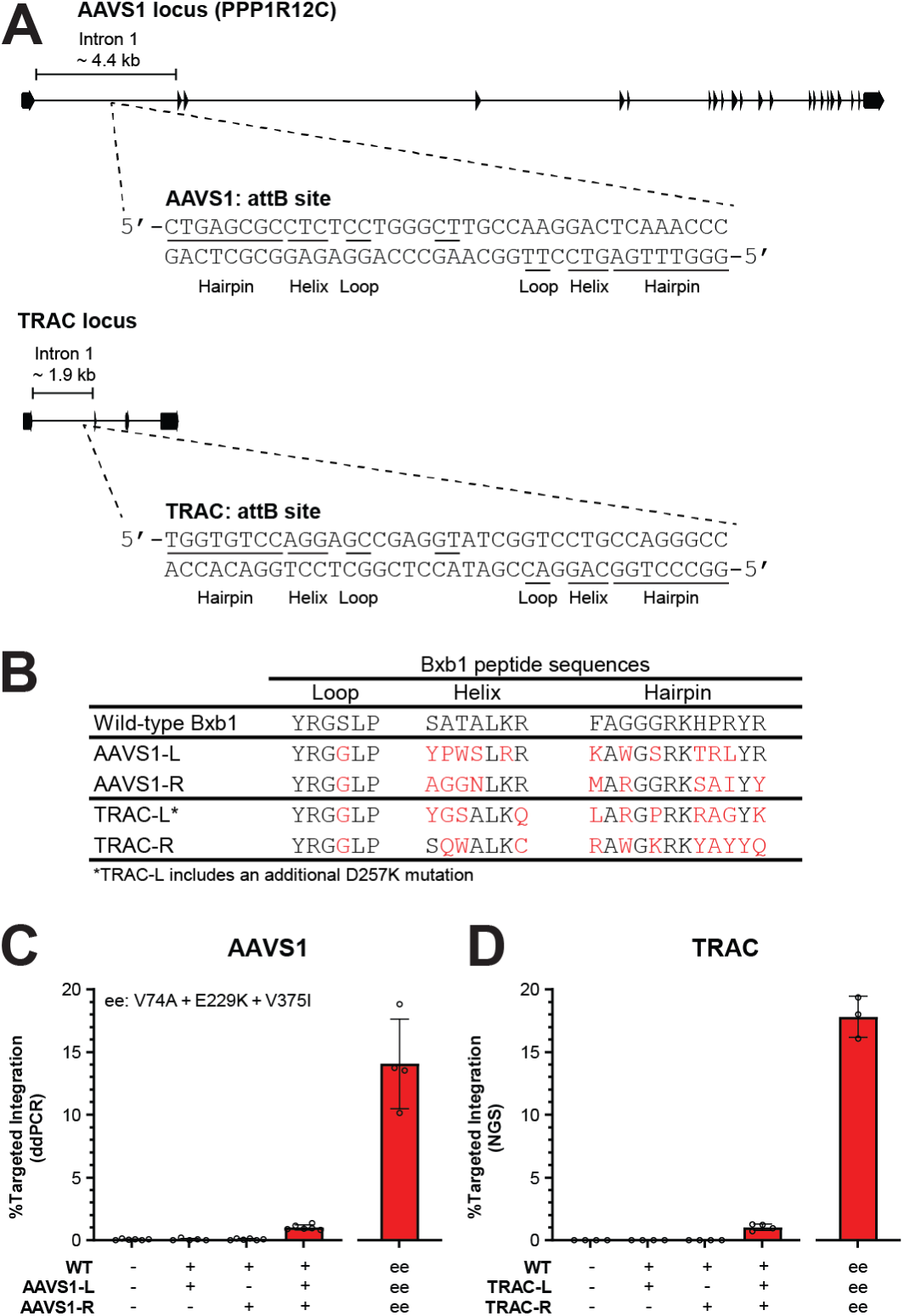
Retargeting Bxb1 to the human TRAC and AAVS1 loci. **a.** Sequence of the attB sites recognized by the TRAC and AAVS1 Bxb1 variants shown in panel b. Both attB target sites are within the first intron of each gene. **b.** Bxb1 peptide sequences of evolved Bxb1 variants for each of the two TRAC and AAVS1 half-sites generated by the strategy shown in Figure 3a. **c.** Results from a digital droplet PCR-based assay demonstrating TI at the depicted AAVS1 site. See Extended Data Fig. 8 for additional details. **d.** Results from our PCR-based NGS assay demonstrating TI at the depicted TRAC site. Data are presented as the mean ± s.d. from four biological replicates for TRAC, and six biological replicates for AAVS1 respectively. Plotted data are provided in the Source Data file.

### Optimization of TRAC-targeted MINT constructs

While we were able to reprogram Bxb1 to target two clinically relevant sites in human cells, the efficiencies that we achieved are not high enough for many therapeutic applications. In order to improve the efficiency of our fully reprogrammed Bxb1 variants that target the human TRAC locus, we investigated two separate approaches: a first approach that involved fusing engineered ZF arrays to our Bxb1 variants and a second approach that involved adding additional mutations that have been shown to increase the activity of wild-type Bxb1 for its natural target site^39^.

Molecular modeling efforts indicated that fusing ZF arrays to the N-terminus of our MINT constructs would likely cause numerous severe steric clashes, so we focused on fusing ZF arrays to the C-terminus of our MINT constructs. Our modeling efforts also indicated which strand of DNA we should target with our ZF arrays such that the N-terminus of each array sits as close as possible to the C-terminus of the cognate Bxb1 variant when both are bound to their intended target sites. In order to evaluate this approach, we designed and tested a panel of linkers and ZF arrays to identify the optimal distance between the 3’ edge of the ZF target site and the 5’ edge of the cognate MINT construct target site. We observed activity improvements when ZF arrays were fused to either the left or right MINT construct and observed the largest improvements when the separation between binding sites was between 4 and 10 bp (**Figure 4a**; Extended Data Fig. 9a). Combining left and right MINT constructs both bearing a fused ZF array further improved performance, yielding more than a 10-fold improvement relative to the same MINT construct without fused ZFs (**Figure 4b**). Encouraged by this result, we next investigated the genome-wide specificity of our TRAC constructs with and without the best performing combination of ZF arrays. We observed a selective increase in signal for the on-target with addition of the ZF arrays with the on-target having 9-fold higher signal than candidate OT1 with the ZF arrays compared to 1.5-fold lower signal than candidate OT1 without these ZF arrays (**Figure 4c)**.

**Figure 4.**
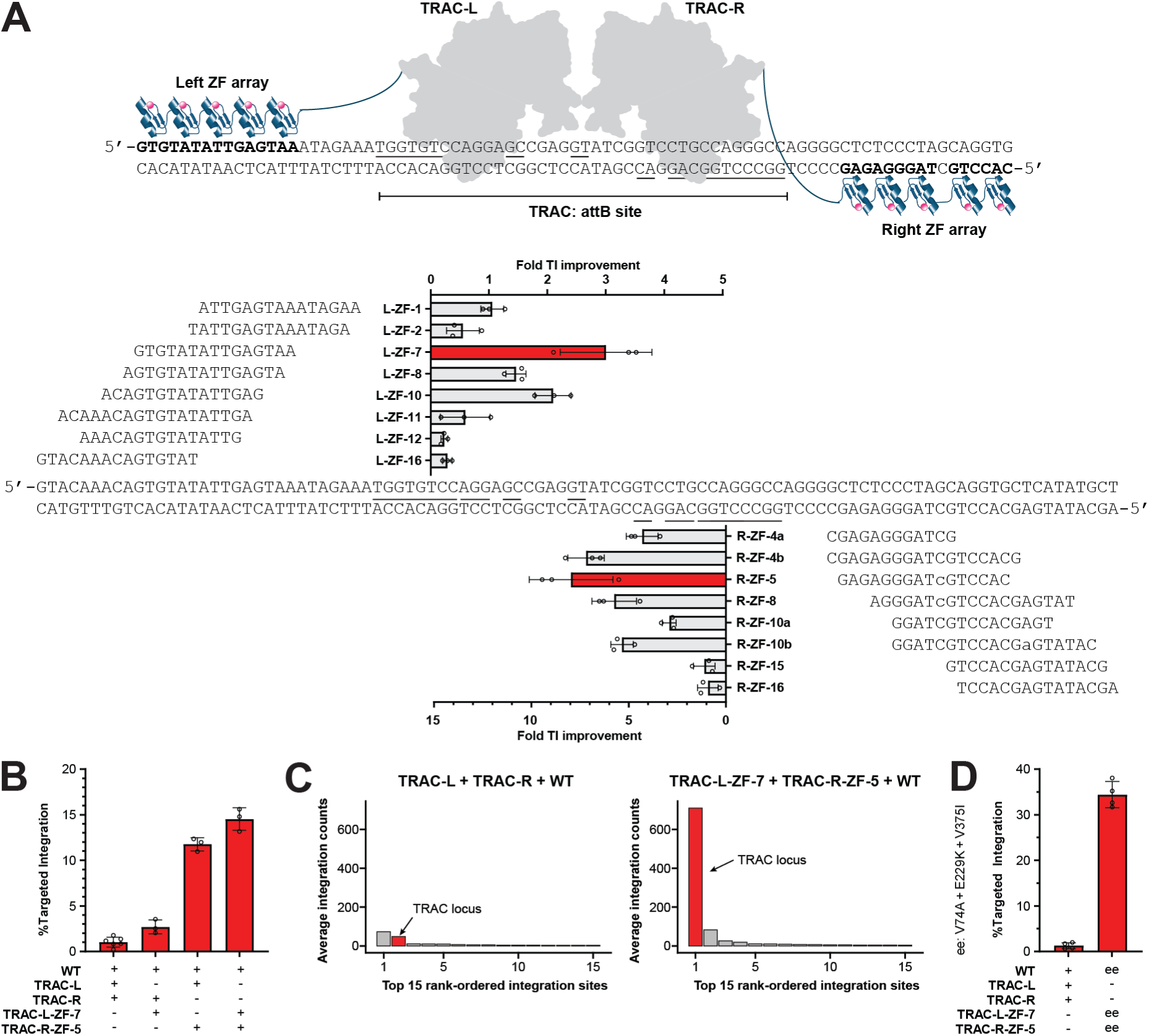
Optimization of TRAC constructs. **a.** Illustration of ZF-fused MINT constructs targeting the human TRAC locus. Eight different ZF arrays were tested for both TRAC-L and TRAC-R. Shown are the respective DNA binding sites and the fold TI improvement when tested by themselves (e.g. L-ZF-1-TRAC-L + TRAC-R + WT). **b.** Demonstration that the best performing constructs from panel a can be paired to further improve efficiency. **c.** Results from an unbiased genome-wide specificity assay demonstrating the selective increase ZF-fusion adds to the on-target integration signal for TRAC MINT constructs. The top 15 candidate integration sites are shown (out of total 85 sites for TRAC constructs and 94 for ZF-fused TRAC constructs) with the entire datasets provided in Supplementary Tables 14 and 15 and corresponding control datasets in Supplementary Tables 16 and 17. Average integration counts were determined using the average deduplicated read numbers across 2 replicates and 2 samples each using primers that amplify the attL and attR plasmid-genome junctions. **d.** Demonstration that activity-increasing Bxb1 variants such as eeBxb1 can be combined with ZF-fused MINT constructs. Data in panels a-c derived from a PCR-based NGS assay and presented as the mean ± s.d. from three to four biological replicates. Plotted data are provided in the Source Data file.

As an orthogonal method to improve integration activity, we added the three mutations utilized in eeBxb1 (V74A, E229K, and V375I)^39^ that have been shown to increase the activity of wild-type Bxb1 for its target site. The addition of these three mutations increased the activity of our AAVS1 MINT constructs by 14-fold (**Figure 3c**) and increased the activity of our TRAC MINT constructs by 17-fold (**Figure 3d**). Finally, we combined both approaches with our TRAC MINT constructs and achieved 35% TI into the TRAC locus in human K562 cells (**Figure 4d**) although the activity increasing mutants did also increase off-target activity (Extended Data Fig. 9b and 9c, Supplementary Tables 18-20). To characterize the nature of the integration, we performed single cell cloning in K562 cells and confirmed full-length TI (Extended Data Fig. 10).

### Performance of MINT constructs in primary human T cells

Encouraged by the high TI levels at the TRAC locus observed in human K562 cells, we decided to test our best performing MINT constructs in the more therapeutically relevant setting of primary human T cells. Since plasmid DNA has been shown to be poorly tolerated by primary human cells^53,54^, we delivered the Bxb1 variants as mRNAs and delivered the donor construct as a nanoplasmid (circular double-stranded DNA lacking a bacterial original of replication and a bacterial antibiotic resistance marker). In order to mimic a therapeutic targeted insertion event, we added a GFP expression cassette to the donor and monitored integration stability and GFP expression. Primary human CD4+/CD8+ T cells were activated and the Bxb1 variants and the donor were delivered to the cells by nucleofection 3 days post-activation. After nucleofection, T cells were incubated for 14 days to allow the GFP signal from unintegrated donor to declined and then TI was characterized by DNA sequencing and GFP expression was analyzed by flow cytometry (**Figure 5a**). MINT construct treatment did not markedly affect T cell growth after recovery from the transfection and T cell markers were stably maintained over the course of the experiments at comparable levels to untreated control cells. Integration activity with the unoptimized TRAC MINT constructs was noticeably higher in T cells than in K562 cells perhaps due to the higher levels of TRAC expression and greater chromosomal accessibility of the TRAC locus in T cells. As in K562 cells, ZF fusions also markedly increased activity in T cells and we achieved 10% TI and 11% stable GFP expression. MINT constructs with both ZF fusions and the activity-increasing mutations yielded 15% TI (**Figure 5b**) and 26% stable GFP expression 14 days post nucleofection (**Figure 5c**).

**Figure 5.**
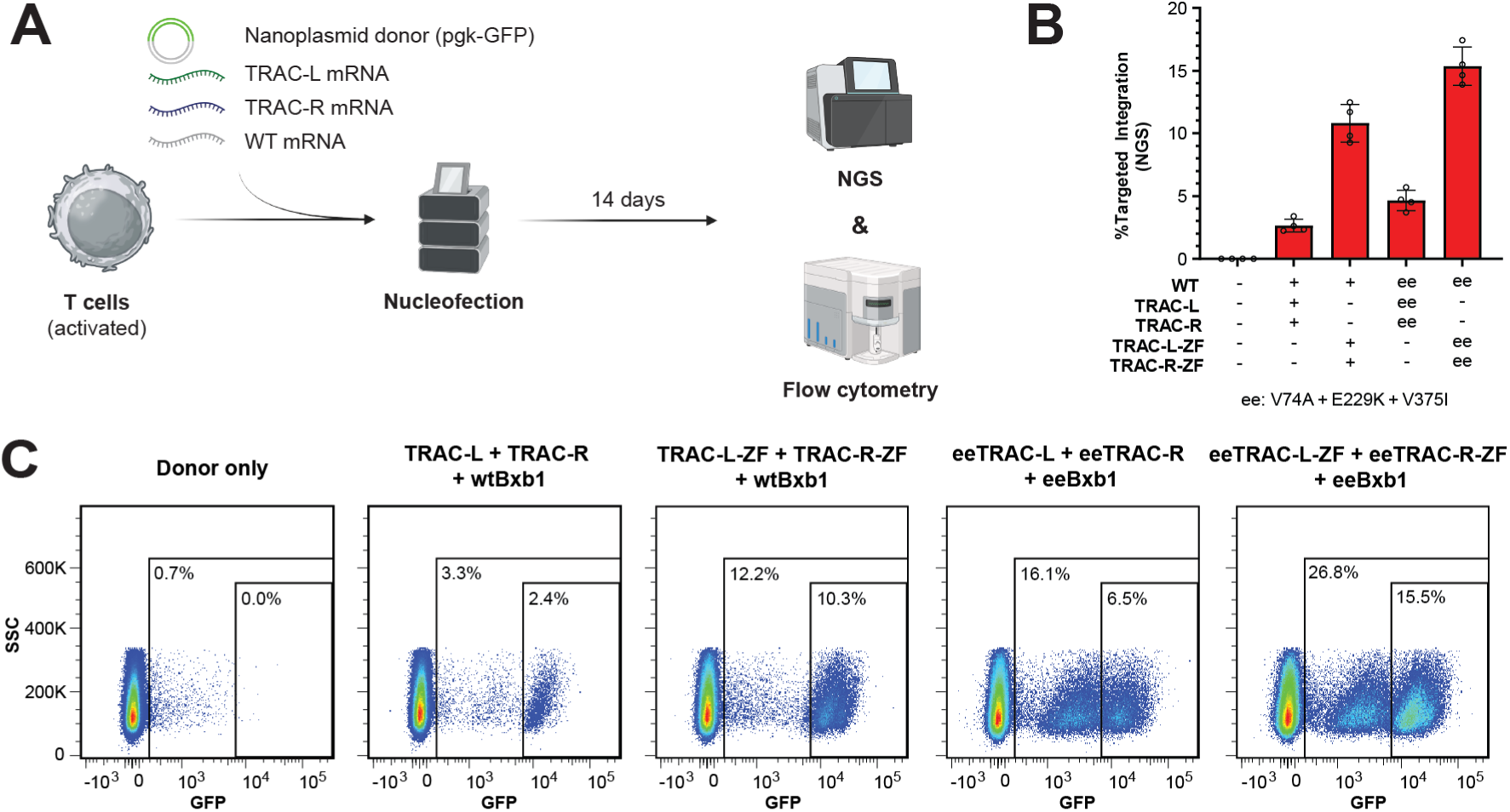
Testing MINT TRAC constructs in primary human T cells. **a.** Experimental procedure of T cell nucleofection and TI quantification. **b.** TI percentage determined by next generation sequencing analysis on DNA isolated 14 days after nucleofection with the indicated constructs. **a** was created with BioRender.com. **b.** Flow cytometric analysis of GFP expression 14 days after nucleofection. The percentage of the GFP-positive cells is indicated. TRAC-L-ZF-7 and TRAC-R-ZF-5 from Figure 4 were used in all T cell nucleofections. Next generation sequencing data are presented as the mean ± s.d. from 4 biological replicates. The flow cytometry plots show individual representative samples. Plotted data are provided in the Source Data file.

We observed subpopulations of cells exhibiting distinct levels of GFP expression, with the high GFP expressing subpopulations more closely mirroring the TI results obtained by NGS (**Figure 5b,c**). To further characterize the subpopulations, we sorted T cells based on GFP expression levels and analyzed TI levels in the subpopulations. The result shows that cells with TI into the TRAC locus are located in the high GFP expressing subpopulations, and that TI percentages in the high GFP expressing subpopulations in MINT-ZF treated cells reach up to 71%, indicating a high frequency of biallelic integration in the highest GFP expressing subpopulation (Extended Data Fig. 11).

## Discussion and future directions

In this work we have demonstrated the direct reprogramming of a site-specific LSR which has been a long-standing challenge for genome engineering^16^. We achieved this through a combination of AI-assisted structural modeling, directed evolution, and screening in human cells that should be applicable to a wide range of site-specific DNA binding proteins. We were able to use this approach to generate numerous reprogrammed Bxb1 variants capable of integrating synthetic DNA into therapeutically relevant endogenous loci in the human genome with efficiencies of up to 35%. To demonstrate that the MINT platform is compatible with AAV-mediated donor delivery, we successfully tested the use of both single-stranded AAV and self-complementary AAV (Extended Data Fig. 1c,d). By combining the MINT platform with known activity-increasing Bxb1 mutants, and fusing ZF proteins to engineered Bxb1 variants, we were able to achieve 15% TI of a functional donor construct into human T cells. We also observed that on-target activity gains from ZF fusions translated better to human T cells than mutations intended to make Bxb1 more active in general.

In contrast to methods that use DNA or RNA replication, such as homology-dependent repair (HDR) or target-primed reverse transcription (TPRT), integrase techniques that use a cut-and-paste mechanism have no inherent limit to the size of the DNA fragment that can be integrated and remove transgene mutation risks caused by use of polymerases. This is not only beneficial for the integration of synthetic DNA into AAVS1 and TRAC, as demonstrated in this study, but also for targeting the first intron of a mutated gene to integrate the correct copy of the corresponding coding sequence linked to a splice acceptor. Furthermore, targeting endogenous attP-like sites in addition to attB-like sites would support other genomic rearrangements such as deletions and inversions. Additionally, the ability to target two different endogenous loci in close proximity would enable recombinase mediated cassette exchange (RMCE) at endogenous loci which could replace an entire genomic locus with a linear synthetic DNA donor construct. This would enable unprecedented flexibility in types of genomic alterations that can be generated^55^. We were able to develop MINT constructs capable of targeting a site within the ∼4 kb human AAVS1 locus and a ∼2 kb intron of the human TRAC locus which demonstrates a high targeting density that would support RMCE at an endogenous locus.

Our results also provide insights into how serine integrases interact with their DNA target sites. Our structural model suggests that residues 231-237 of Bxb1 interact with DNA in a manner reminiscent of how the recognition helix of a ZF interacts with DNA. To our surprise, we observed similar patterns of amino acids at specific positions of the Bxb1 recognition helix that have been observed in ZF engineering efforts (Supplementary Fig. 1). In contrast, the hairpin region appears to use a distinctly different mechanism to recognize DNA. Through an analysis of Bxb1 targets in the human genome, we observed that the hairpin region of wild-type Bxb1 can recognize at least four distinct DNA motifs implying that the hairpin region can either adopt multiple conformations or bind using multiple DNA docking geometries (Extended Data Fig. 4c,d). This is consistent with the observation that the ZD domain of wild-type Bxb1 can specifically recognize two completely different sequences in the left and right half-sites of the natural attB target. The loop region appears to recognize DNA using yet another mechanism. The fact that a single mutation, S157G, can dramatically relax specificity implies that the loop region may act more as a switch that reduces integration activity when the preferred sequence is not detected rather than the more standard model of interactions that increase binding affinity when the preferred sequence is present.

We envision that our MINT platform will also accelerate efforts in other research areas where Bxb1 has been successfully used, such as for metabolite pathway assembly^56^ and various agrobiotechnology applications in model plants^57^ and crops^58,59^. We believe the modularity and efficiency of our approach will make it an ideal choice for applications where error-free integration of large synthetic DNA constructs is required.

In this study we established a method for retargeting Bxb1 that will likely enable the reprogramming of other serine integrases including those that have recently been described^38,60^ (Supplementary Fig. 2). Since the RD and ZD domains of Bxb1 behave in a modular fashion it is likely that RD and ZD domains from different LSRs can be combined to further expand the ability to target desired genomic loci. Ultimately, our goal is to create an archive of pre-characterized integrase helix and hairpin variants. Such a pre-characterized archive would allow the *in silico* design of Bxb1 variants to target any sequence of interest by simply combining the appropriate helix variant with the appropriate hairpin variant without the need for custom directed evolution experiments. This work provides an initial archive of 26 hairpin variants and 24 helix variants that are active in human cells (Supplementary Fig. 3,4; Supplementary Tables 22-24) that can be used in conjunction with activity-increasing mutations and ZF fusions to target desired genomic loci for therapeutic applications and beyond.

## Supporting information

MINT_Sequence_Information

MINT_Source_Data

MINT_Supplementary_Tables

## Data availability

Amino acid sequences and DNA sequences of constructs used in this study are provided as supplementary information. Illumina sequencing data underlying all experiments will be deposited in the NCBI Sequence Read Archive before publication. Source data are provided with this paper.

## Code availability

All custom python scripts will be made available on GitHub after acceptance of this manuscript at a peer-reviewed scientific journal.

## Acknowledgements

We thank Gustavo de Alencastro for assistance with AAV construct design, as well as Sumita Bhardwaj, Tammy Chen, Alicia Goodwin, Emma Petrouski, Sanjna Sridhar, and Hung Tran for assistance with AAV production. We thank Trevor Collingwood for scientific discussions. We also thank Ed Rebar, Jon Melnick, Deepak Patil, and Charles Paine for contributions to the early development of ZF-targeted Serine Recombinases. And we thank Sandy Macrae, Adrian Woolfson, and Jason Fontenot for their relentless support of this project.

## Author contributions

F.F., S.A.-F., J.E.D., L.L, N.J.S., L.R., N.A.S., R.M., I.T., B.N.K., M.Q, P.L., A.R. and J.C.M. designed experiments. F.F., S.A-F., J.E.D., L.L., N.J.S., L.R., D.F.X., N.N., N.A.S., R.M., I.T., Y.Z., L.N.T., S.J.H., B.N.K., S.L., B.B., E.T., M.Q., V.V., A.C., A.N., and P.L. performed experiments. F.F., S.A.-F., J.E.D., L.L., N.J.S., L.R., D.F.X., N.N., N.A.S., R.M., I.T., Y.Z., L.N.T., S.J.H., B.N.K., S.L., B.B., E.T., M.Q., V.V., A.N., P.L., A.R. and J.C.M. performed data analysis. F.F., S.A.-F., J.E.D., L.L., S.J.H., P.L., A.R., D.E.P., G.D.D., and J.C.M. supervised experiments. J.E.D., Y.R.B., D.A.S., and J.C.M. developed custom computer code. F.F., S.A.-F., and J.C.M wrote the manuscript with input from all authors.

Material requests and correspondence to F. Fauser and J.C. Miller.

## Methods

### Cloning of expression constructs and donors used in mammalian cells

Most constructs were cloned using NEBuilder® HiFi DNA Assembly (NEB, Catalog #E2621X) or Q5® Site-Directed Mutagenesis (NEB, Catalog #E0554S), or synthesized and cloned by Twist Biosciences utilizing their clonal gene services. The gBlocks or eBlock DNA fragments used for constructs assembly were synthesized at IDT (Integrated DNA Technologies). DNA sequences for all constructs can be found in supporting sequence information. All constructs were sequence confirmed using Sanger sequencing services provided by Elim Biopharm Inc., and whole plasmid sequences were verified using either the Nextera XT DNA library prep kit (Illumina, Catalog #FC-131-1096) or whole plasmid sequencing services (Plasmidsaurus Inc.; Elim Biopharma Inc.).

The ZF constructs have the following amino acid sequences where recognition helix sequences are underlined and structured basepair skipping linkers are in italics:

### High throughput Bxb1 variants assembly

The Bxb1 variant gene fragments (bases 460 – 1087, corresponding amino acids 144 – 362 with loop, helix and hairpin regions included) were synthesized as eBlocks (Integrated DNA Technologies) and assembled to full length fusion expression cassette through 2-step PCR. In the first step PCR, the eBlocks were amplified with overlapping gene fragments DF148 and DF164 with AccuPrime *Pfx* SuperMix (Invitrogen, Catalog #12344040) and the following thermocycler conditions: initial melt of 95 °C for 3 min; 15 cycles of 95 °C for 30 s, 68 ° for 30 s, and 68 °C for 2 min 30 s; followed by a final extension at 68 °C for 5 min; hold at 4 °C. The full-length expression cassette PCR products were then amplified with forward primer 5’-GCAGAGCTCTCTGGCTAACTAGAG-3’ and reverse primer 5’-TTTTTTTTTTTTTTTTTTTTTTTTTTTTTTTTTTTTTTTTTTTTTTTTTTTTTTTTTTTTTAGACAGGCGGGGAGGCGGC-3’ and the following thermocycler conditions: initial melt of 95 °C for 3 min; 30 cycles of 95 °C for 30 s, 68 ° for 30 s, and 68 °C for 2 min 40 s; followed by a final extension at 68 °C for 5 min; hold at 4 °C. The final PCR products were examined on a gel and purified with AMPure XP beads (Beckman# A63881). The purified PCR products were used for mRNA production.

### mRNA production

The mRNA was prepared with purified PCR products via the mMESSAGE mMACHINE T7 Ultra kit (ThermoFisher, Catalog #AM1345) following the manufacturer’s instructions. Synthesized mRNA was purified with RNA cleanXP beads (Beckman, Catalog #A63987) and quantified with Quant-IT kit (Invitrogen, Catalog #Q10213).

### AAV production

AAV genome plasmids containing the relevant attP sequences for Bxb1-mediated recombination, flanked by inverted repeat terminal sequences (ITR) for AAV packaging, were cloned using NEBuilder® HiFi DNA Assembly (NEB, Catalog #E2621X). ITRs were mutated to produce scAAV^61^. Bovine growth hormone (bGH) polyA termination sequence was inserted upstream of the 3’ ITR to aid transcription termination as well as AAV titration. All constructs were sequence confirmed using Sanger sequencing services provided by Sequetech Corporation. The whole plasmid sequences were verified using Nextera XT DNA library prep kit (Illumina, Catalog #FC-131-1096) or whole plasmid sequencing services provided by Plasmidsaurus Inc. to ensure ITRs are intact prior to AAV production.

The recombinant AAV vectors (rAAV) were produced by triple transfection of suspension human embryonic kidney (HEK) cells in 400 ml flasks, then purified by cesium chloride density gradient centrifugation and dialysis. The rAAV genome was packaged into AAV-DJ (synthetic chimeric capsid of AAV 2/8/9) and titrated using bGH PolyA qPCR assay on the Quantstudio™ 3 Real-time PCR system.

### Directed evolution system

#### Preparation of Integrase Mutant libraries

As a starting point for engineering of Bxb1 variants in E. coli, we cloned a codon optimized ORF into plasmid pBxb1, which contains a pBR322 origin of replication as well as the L-rhamnose inducible pRhaBAD promoter. Libraries were prepared using inverse PCR and primers encoding degenerate bases using an NNK degeneracy scheme. In order to diversify the “loop” submotif, primers were designed to target residues 154-159. In order to diversify the “helix” submotif, primers were designed to target residues 231-234, and residues 236-237. In order to diversify the “Beta-hairpin” submotif, primers were designed to target residues 314, 316, 318, 321, 323, and 325 using an NNK randomization scheme, while residue 322 was randomized using an SSK randomization scheme. Briefly, iPCR reactions were set up in 50 uL volumes, using 1x KOD ONE Master mix, 0.5uM forward primer, 0.5 uM reverse primer, and 10 ng of template plasmid DNA. Following purification of the PCR products via silica column (Qiagen PCR cleanup kit), and elution in 50 uL of buffer EB, compatible overhangs for ligation were generated by digestion of PCR products using 5 units each of BsaI (NEB, R3733S) and DpnI (NEB, R0176S) in 60 uL of 1x CutSmart buffer. The resulting digest products were then ligated at 10 ng/uL in 1x T4 Ligase buffer, at 8 U/uL T4 Ligase, for 1 hour at room temperature. Following ligation, DNA products were purified using the Qiagen PCR cleanup kit and eluted in 50 uL of buffer EB. The resulting ligated products were used to transform 50 uL of electrocompetent cells (NEB 10Beta, C3020K; or ThermoFisher OneShot Top10, C404052) using a BMX electroporator and a 96-well cuvette (BTX, 45-0450-M) using the manufacturer’s protocol and allowed to recover for 1 hour at 37 °C in 1ml total volume of SOC media. The resulting transformations yielded libraries in the order of 4e8 to 5e9 CFUs. After recovery, cells were transferred to 1 L of LB media containing 34 ug/mL chloramphenicol and grown overnight with shaking at 37 °C. The resulting culture was harvested, and plasmids were purified using a Qiagen Plasmid Plus Giga Kit.

#### Preparation of selection cassette plasmids

Selection plasmid pTarget was based on a p15A origin of replication and contained a spectinomycin resistance cassette. The selection cassette contained a Tet promoter upstream of an open reading frame encoding a gentamicin resistance gene (aacC1) disrupted by a “stuffer sequence”. The stuffer sequence was flanked with extra sequence upstream of attB and downstream of attP to enable changing of reading frames if attL recombination produced an unviable peptide stuffer sequence. The attB and attP sequences were modified such that recombination by an active integrase would leave behind an attL sequence encoding an in-frame peptide insertion, giving rise to an active aacC1. Each selection cassette was generated by the cloning of two ∼1050 bp DNA fragments into a linearized pTarget backbone plasmid using the NEB HiFi assembly master mix, according to the manufacturer’s instructions. Sequence-confirmed plasmids were either prepared by Mini- or Midi-Kits (Qiagen) according to the manufacturer’s instructions.

#### Selection of Bxb1 variants by aacC1 reassembly

Typical selection reactions used 1ug of Bxb1 library, and 1ug selection cassette plasmid. OneShot Top10 electrocompetent cells were mixed with library and selection plasmids before being transferred to a 96-well cuvette, and electroporated as above. After recovery, cells were transferred to 4 mL of LB containing 34 ug/mL chloramphenicol, 50 ug/mL spectinomycin and 0.2% L-rhamnose, then the cultures were shaken at 37 °C for 16 h to allow recombination to take place. For selections using liquid broth, cells were spun down and resuspended in 50 mL Super Broth containing 34 ug/mL chloramphenicol and 20 ug/mL gentamicin. After overnight culture, 1 mL of each culture was used to extract plasmid DNA using the Qiagen Miniprep kit. For selections using agar plates, cells were instead spun down and resuspended in 1 mL of Superbroth and plated on 25×25 cm LB agar plates containing 34 ug/mL chloramphenicol and 20 ug/mL gentamicin. After overnight incubation at 37 °C, colonies were scraped from plates, resuspended in 10 mL LB broth, and a 500 uL volume was removed and used in plasmid purification using Qiagen Miniprep kits.

#### NGS sequencing of Bxb1 variant submotifs post-selection

Samples were prepared for sequencing using Illumina MiSeq or NextSeq using an integrase-adaptor hybrid primer pair (PCR1) followed by an Illumina-adaptor specific primer pair (PCR2). Briefly, 50uL reactions were prepared using 1x Q5 Hotstart Master mix (NEB M0494S), 0.5uM Forward primer, 0.5uM Reverse primer, and 50ng of plasmid DNA. PCR was performed using the following cycling conditions, 98 °C 30 s, then 15 cycles of 98 °C 5 s, 68 °C 7 s, 72 °C 10 s, followed by a final extension step at 72 °C for 30s. The resulting PCR product was used as template for PCR2. Briefly, 50 uL reactions were prepared using 1x KOD ONE polymerase master mix (Millipore-Sigma), 0.5 uM Adapter primer 1, 0.5 uM Adapter primer 2, and 1 uL of PCR1 as template. PCR was performed using the following cycling conditions 98 °C 30 s, then 13 cycles of 98 °C 5s, 60 °C 7 s, 72 °C 5 s followed by a final extension step at 72 °C for 30 s. The resulting PCR products were column purified using a Qiagen PCR cleanup kit, and samples were sequenced using standard Illumina Kits for MiSeq or NextSeq.

#### Molecular specificity assays

We built a plasmid reporter to assay the molecular specificity of novel integrase submotifs. Three versions of the pBxb1 plasmids were used as starting point to avoid background activity of the wild-type integrase during cloning. By placing an extra adenine residue within the loop, helix, or hairpin submotif, we created pBxb1 variants where Bxb1 was inactivated by a frameshift-inducing mutation but could be rescued by subsequent mutagenesis. Recombination cassettes from the pTarget selection plasmids were amplified by PCR and cloned into the pBxb1 plasmids by Gibson assembly upstream of the Bxb1 ORF. To assay the specificity of selected loops, a single reverse primer and different forward primers which encoded the new loop residues were used in iPCR using the small library of loop target plasmids (16 targets). A similar procedure was used to generate helix mutants using the small library of helix target plasmids (64 targets).

To generate a library of ZD hairpin targets, we amplified and cloned an oligo pool (Twist Biosciences) which comprised all possible attB and attP sites where positions −18 to −13 have been randomized into our pBxb1 plasmid, downstream of the integrase ORF. Such a library ensures that the ZD hairpin targets on attBL and attPL are identical, and the targets on attBR and attPR are inverted repeats of the left side targets. Mutant hairpins are generated by two different iPCR primers, which together encode the novel hairpin sequence.

iPCR reactions were carried out in 50 uL volumes, with 1 x KOD ONE master mix, 0.5uM forward primer, 0.5uM reverse primer, and 20 ng of a plasmid pool containing the relevant recombination cassettes. Thermocycling was carried out using an initial denaturation step of 98 °C for 30s, followed by 35 cycles of 98 °C 10s, 60 °C 5s, 68 °C 30s. PCR amplicons were purified using AMPure XP beads (Beckman) and eluted in 40 uL EB buffer (Qiagen). 30 uL of the purified PCR amplicons were then digested and ligated in a 50 uL one-pot reaction containing 1x T4 ligase buffer (NEB), 20U DpnI (NEB), 20 U BsaI-HFv2 (NEB), and 400 U of Salt-T4 DNA Ligase (NEB). All reactions were then incubated at 37 °C 30 minutes, 20 °C 30 minutes, 37 °C for 30 minutes. 20 uL of thawed chemically competent NEB 5alpha cells were added 2 uL of the digested/ligated amplicons, and cells were transformed according to the manufacturer’s instructions. After recovery, cells were added to 800 uL of media containing 34ug/mL chloramphenicol and 0.2% L-rhamnose (w/v). The resulting culture was incubated for 16 hours at 37 °C with shaking in 96-well deep-well plates, then harvested by centrifugation. Plasmid DNA was extracted using a Qiaprep 96 Turbo Miniprep kit.

To assess the specificity of each clone, primers which flank the resulting attL sequence and contain Illumina adaptor sequences were used to amplify the products of recombination. Briefly, PCR reactions were carried out in 20uL volumes, using 1 x Hotstart Taq master mix (NEB, M0496S), attB_MiSeq forward primer, and attP_Miseq reverse primer, using 1uL of purified plasmid as template. Thermocycling was carried out as follows 98 °C 30s; 25 cycles of [98 °C 5s, 53 °C 10s, 72 °C 5s] Final extension 72 °C 30seconds. A second PCR to install sequencing barcodes was carried out using specific barcoding primers, with the recombination products from each clone being represented by a unique combination of forward and reverse barcoding primers. PCR reactions were carried out in 20 uL volumes, using 1 x KOD ONE master mix, 0.5 uM forward primer, 0.5 uM reverse primer, and 1 uL of the previous PCR product. Thermocycling was carried out as follows 98C 30s; 12 cycles of [98 °C 5s, 60 °C 5s, 68 °C 3s]. The resulting PCR products were column purified using Qiagen PCR cleanup kits, then sequenced using standard Illumina kits for NextSeq or MiSeq.

### General mammalian cell culture conditions

K562 cells (ATCC, CCL243) were cultured using RPMI-1640 growth medium supplemented with 10% FBS (Fetal Bovine Serum) and 1x PSG (Penicillin-Streptomycin-Gentamycin, Gibco, 10378-016) and maintained at 37 °C with 5% CO2.

CD4+/CD8+ primary human T cells were purified from healthy donor leukopaks. Primary human T cells were cultured in X-Vivo 15 (Lonza, BP04-744Q) supplemented with HEPES (Gibco, 15630-080), GlutaMAX (Gibco, 35050-061), Sodium Pyruvate (Gibco, 11360-070), MEM vitamins (Corning 25-020-c1), human AB serum (Valley BioMedical, HS1017HI) & IL-2 (Invitrogen CTP0023).

### K562 tissue culture nucleofection protocol and genomic DNA preparation

Expression constructs were routinely dosed as plasmid DNA (pDNA) in K562 cells. K562 cells were electroporated with pDNA using the SF cell line 96-well Nucleofector kit (Lonza, Catalog#V4SC-2960) or SF Cell Line 384-well Nucleofector Kit (Lonza, Catalog#V5SC-2010), using manufacturer’s protocol. Prior to electroporation, K562 cells were centrifuged at ∼300 x g for 5 min, and washed with 1x DPBS (Corning, Catalog#21-031-CV). For 96-well nucleofection, cells were resuspended at 2e5 cells per 12 µl of supplemented SF cell line 96-well Nucleofector solution. 12 µl of cells were mixed with 8 µl of pDNA and transferred to the Lonza Nucleocuvette plate. Nucleofector program 96-FF-120 was used to electroporate K562 cells with the pDNA mix on the Amaxa Nucleofector 96-well Shuttle System (Lonza). After electroporation, cells were incubated for 10 min at room temperature and transferred to a 96-well tissue culture plate containing 180 µl of complete medium (prewarmed to 37 °C).

For 384-well nucleofection, cells were resuspended at 1e5 cells per 14 µl of supplemented SF cell line 384-well Nucleofector solution. 14 µl of cells were mixed with 6 µl of pDNA including 1400ng donor, 66ng of wt, 66ng of left and 66ng right bxb1 variants or bxb1 variant-ZFP fusion DNA and transferred to the Lonza Nucleocuvette plate. Nucleofector program FF/120/DA was used to electroporate K562 cells with the pDNA mix on the Amaxa HT Nucleofector System (Lonza, AAU-1001). After electroporation, cells were incubated for 10 min at room temperature and transferred to a 384-well tissue culture plate containing 60 µl of complete medium (prewarmed to 37 °C). K562 cells were incubated for ∼72 h and then harvested for quantification of editing events.

### HepG2 tissue culture nucleofection protocol and genomic DNA preparation

HepG2 cells were electroporated with pDNA using the SF cell line 96-well Nucleofector kit (Lonza, Catalog #V4SC-2960) using manufacturer’s protocol. 1000 ng of Bxb1 or GFP control along with 5000 ng of plasmid donor was used. Prior to electroporation, HepG2 cells were centrifuged at ∼200 x g for 10 min, and washed with 1x DPBS (Corning, Catalog #21-031-CV).

For 96-well nucleofection, cells were resuspended at 2e5 cells per 20 µl of SF Nucleofector Solution with supplement. 16.5 µl of cells were mixed with 3.5 µl of pDNA, then transferred to the Lonza 96-well Shuttle plate. Nucleofector program 96-EH-100 was used to electroporate HepG2 cells with the pDNA mix on the Amaxa Nucleofector 96-well Shuttle System (Lonza). After electroporation, cells were incubated for 10 min at room temperature, 180 ul of growth medium (MEM plus 10% FBS) were added per well and cells transferred to a 48-well tissue culture plate containing 300 µl medium (prewarmed to 37 °C). Electroporated HepG2 cells were placed in a 30 °C incubator for 24 hours and then transferred to a 37 °C incubator. HepG2 cells were incubated for ∼72 h and then harvested for quantification of editing events.

### Digital Droplet PCR quantification of targeted integration at AAVS1 target site

For AAVS1 target integration ddPCR quantification, 2 TI probes (targeting attL and attR) and 2 reference probes were designed. The target integration probes were designed targeting either attL or attR region which would not be detected in non-transfected cells or plasmid donors. The reference probes were designed within a 10 kb region of the AAVS1 attB target site to mitigate the risk of copy number difference due to abnormal karyotype in K562 cells. The probes and primers were designed with PrimerPlus3 (https://www.primer3plus.com/) based on the recommendation by BioRad manual and were synthesized at IDT (Integrated DNA Technologies, Inc.). The primers and probes were tested in duplex format at different annealing temperatures and with synthetic eblock mixes, non-transfected cell lysate and NTC. The attL and reference2 probes and primers and 57.1°C were selected for AAVS1 target site ddPCR quantification.

DNA was extracted with QuickExtract DNA Extraction Solution (Lucigen#QE09050). For each reaction, 50 µl of QuickExtract DNA solution was added to approximately 0.5-1 million pelleted cells, followed by mixing and incubation at 65 °C for 15 min and heat inactivation at 98 °C for 5 min. The cell lysates were mixed by vortexing for 15 seconds before ddPCR. Each ddPCR reaction was prepared and analyzed with a QX200 ddPCR system (Bio-Rad) and ddPCR Supermix for Probes (No dUTP) (Bio-Rad, Catalog #1863024) per Bio-Rad’s standard recommendations. All reactions were mixed to 22 µl including 10 U of HindIII-HF (NEB, Catalog #R3104L) and up to 2 µl of QuickExtract lysates. Forward primer, reverse primer and probe were at a 3.6:3.6:1 ratio. Droplets were generated in the droplet generator per Bio-Rad’s protocol. Thermocycler conditions: 95 °C for 10 min; 40 cycles of 95 °C for 30 s and 57.1 °C for 60 s; 98 °C for 10 min; and hold at 8 °C. QX Manager Software 2.1 Standard Edition (Bio-Rad) was used for QC and the analysis. The thresholds were set manually at 3000 for channel1/FAM and 1000 for channel2/HEX. All final data was exported into Microsoft Excel for further analysis. The target integration ratio was calculated by the equation: TI (%) = 100*CattL/Cref2 (C: volumetric concentration (copies/μl)).

### AAV transduction

For evaluating rAAVs. K562 cells were electroporated with 800ng of Bxb1 pDNA using the SF cell line 96-well Nucleofector kit (Lonza, Catalog#V4SC-2960), using manufacturer’s protocol. K562 cells were centrifuged at ∼300 x g for 5 min, and washed with 1X PBS (Corning, Catalog#21-031-CV). For 96-well nucleofection, cells were resuspended at 2E5 cells per 12 µl of supplemented SF cell line 96-well Nucleofector solution. 12 µl of cells were mixed with 8 µl of Bxb1 pDNA and transferred to the Lonza Nucleocuvette plate. Nucleofector program 96-FF-120 was used to electroporate K562 cells with the pDNA mix on the Amaxa Nucleofector 96-well Shuttle System (Lonza). After electroporation, cells were incubated for 10 min at room temperature and transferred to a 96-well tissue culture plate containing 180 µl of complete medium (prewarmed to 37 °C). 30 mins post-electroporation, rAAV constructs were dosed at MOI (multiplicity of infections) at 500,000 vg/cell. rAAV donor only control wells were included in parallel. K562 cells were incubated for ∼72 h and then harvested for quantification of editing and circularization events. A PCR-based NGS assay was used to measure TI events.

### K562 landing pad cell line generation

K562 cells were electroporated with a pair of AAVS1 zinc finger nuclease (ZFN) mRNA and an ultramer with the attB sequence (see supplementary sequence information) using the SF cell line 96-well Nucleofector kit (Lonza, Catalog#V4SC-2960), using manufacturer’s protocol. The ZFNs generate a double-strand break in the genome that facilitates integration of the ultramer via homology directed repair through the corresponding homologous ends. K562 cells were incubated for ∼72 h at 37 °C with 5% CO2. One third of the cells were harvested for quantification of bulk integration of the ultramer using a PCR-based NGS assay. 2/3^rd^ of the cells were maintained for diluting to singles. Samples showing ∼10% integration were selected and diluted to singles in a 96-well plate and incubated for ∼1.5 weeks. At the end of ∼1.5 weeks, the plates were examined under a microscope for the growth of single clones. Cells from wells showing single clones were transferred to a 24-well tissue culture plate and transferred to 37 °C with 5% CO2 for 3 days or until 75-80% confluency is reached. 50% of the cells were harvested for quantification of ultramer integration using the PCR-based NGS assay. Cells were maintained in fresh medium until the NGS assay was completed. Since AAVS1 has 3 alleles, a sample showing 34.39% integration at a single allele was selected for further expansion. The other two wild-type alleles showed a 6 bp deletion, but this did not disrupt the performance of the cell line.

### PCR-based NGS assay for targeted integration and indel quantification

3 days post transfection, cells were spun down at ∼500 x g for 5 min. Supernatant was discarded, cells were washed in PBS (Corning, Catalog #21-031-CV), and cells were resuspended in 50 µl of QuickExtract DNA Extraction Solution (Lucigen, Catalog #QE09050). Genomic DNA was extracted by treating the cells to the following protocol: 65 °C for 15 min, 98 °C for 8 min. Target sites were amplified from the genomic DNA using Accuprime HiFi reagents (Invitrogen, Catalog #12346094) and the following PCR conditions: initial melt of 95 °C for 5 min; 30 cycles of 95 °C for 30 s, 55 °C for 30 s and 68 °C for 40 s; and a final extension at 68 °C for 10 min. Primers containing adapters (forward primer adapter: ACACGACGCTCTTCCGATCT; reverse primer adapter: GACGTGTGCTCTTCCGATCT), targeting specific target sites were used at a final concentration of 0.1 µM. Sequences for the primers used can be found in supporting sequence information. The PCR products obtained were then subjected to a second PCR to add Illumina barcodes to the PCR fragments generated in the first PCR. We used Phusion High-Fidelity PCR MasterMix with HF Buffer (NEB, Catalog #M0531L) for the second PCR and used the following PCR conditions, initial melt of 98 °C for 30 s; 12 cycles of 98 °C for 10 s, 60 °C for 30 s and 72 °C for 40 s; and a final extension at 72 °C for 10 min. PCR libraries generated from the second PCR were pooled and purified using QIAquick PCR purification kit (Qiagen, Catalog #28106). Samples were diluted to a final concentration of ∼2 nM after they were quantified using the Qubit dsDNA HS Assay kit (Invitrogen, Catalog #Q33231). The libraries were then run on either an Illumina MiSeq using a standard 300-cycle kit or an Illumina NextSeq 500 or an Illumina NextSeq 2000 using a mid-output 300-cycle kit using standard protocol.

### Pooled screening of attB sites in K562 cells

Extended Data Fig. 7a outlines the experimental screening of Bxb1 variants against plasmid libraries of artificial target sequences in human K562 cells. Target libraries were designed as shown in Extended Data Fig. 7b,c and cloned using oligo pools (Twist Bioscience). The resulting target libraries were then co-transfected with a universal donor and individual Bxb1 variants. Activity was measured using PCR-based NGS assay.

### Genome-wide specificity assay

Our genome-wide specificity assay was adapted from the GUIDE-seq protocol^51^, with modifications designed to measure Bxb1-induced off-target editing events. K562 cells were transfected using conditions similar as described above. Transfections involving integrations with multiple donors with different core dinucleotides were performed separately per donor plasmid and then cells were pooled and expanded for 1 week before being spun down for genomic DNA extraction. Without the DpnI site addition to donor plasmids, the cells were grown out for 3-4 weeks prior to DNA extraction and the DpnI digestion step below was not followed.

Genomic DNA was extracted from K562 cells using Qiagen DNeasy Blood & Tissue kits (Catalog #69504) following the Purification of Total DNA from Animal Blood or Cells Spin-Column Protocol for cultured cells. The optional RNase A incubation was followed for all samples. DNA was eluted in 60 µL Elution Buffer and quantified using the Qubit fluorometer and the Qubit dsDNA HS Assay Kit (Invitrogen, Catalog #Q33231) following the recommended protocol.

Next, 10 uM adapters were prepared by annealing the MiSeq common oligo to each GS_i5 oligo in a 96-well plate format to make a barcoded Y adapter plate. Annealing was performed with 1X oligo annealing buffer (10 mM Tris HCL pH 7.5, 50 mM NaCl, and 0.1 mM EDTA) by following the below thermocycling method: initial melt of 95 °C for 2 min, step-down from 80 °C to 4 °C with −1 °C per cycle and 1 min incubation at each temperature, hold at 4 °C until further use. Adapters were stored at −20 °C, and before use were thawed on ice400 ng (133,000 haploid human genomes) genomic DNA was brought up to 50 uL using 1X IDTE pH 7.5 (IDT, Catalog #11-05-01-05) in each tube of a Covaris 8 microTUBE-130 AFA Fiber H Slit Strip V2 (Covaris, Catalog #520239). Samples were sonicated on a Covaris ME220 using the following settings on a ME220 Rack 8 AFA-TUBE TPX Strip (Covaris, Catalog #PN500609) using the waveguide (Covaris PN 500526): Power 0.0 W, Temperature 19.7 °C, Duration(s) 65.0, Peak Power 40.0, Duty %Factor 10.0, Cycles/Burst 10000, Avg. Power 4.0.

Sheared DNA was purified using 1 volume Ampure XP beads (Beckman Coulter, Catalog #A63880). After beads were added, the solution was mixed and incubated for 5 minutes at room temperature. The mixture was then incubated on a magnet for 5 minutes before the supernatant was removed. 150 uL freshly made 70% ethanol was then used to wash the beads twice, allowing the solution and beads to sit for 30 seconds each time. After the second wash, the beads were dried for 6 minutes before adding 15 uL IDTE pH 7.5 and mixing off the magnet. After 2 minutes the mixture was placed on a magnet and incubated for another 2 minutes. 14.5 uL of the supernatant was collected for the next step.

The reaction was brought up to 50 uL with the addition of CutSmart (final concentration 1X) and 1 uL DpnI (NEB, Catalog #R0176S) and incubated for 1 hour at 37 °C. DNA was purified using 0.8x Ampure XP beads using the same bead clean-up protocol as before. Next, the following End repair and A-tailing mixture was added to each 14.5 uL DNA mixture, while the reaction was kept on ice: 0.5 uL 10 mM dNTP mix (Invitrogen, Catalog #18427013), 2.5 uL 10X T4 DNA Ligase Buffer (Enzymatics, Catalog #B6030), 2 uL End-repair mix (Enzymatics, Catalog #Y9140-LC-L), 2 uL 10X Platinum Taq Polymerase PCR Rxn Buffer (-Mg2 free) (Invitrogen, Catalog #10966034), 0.5 uL Taq DNA Polymerase Recombinant (5u/ uL) (Invitrogen, Catalog #10342020), and 0.5 uL dsH2O to a total of 22.5 uL. The solution was mixed and incubated on a thermocycler with the following program: 12 °C for 15 min, 37 °C for 15 min, 72 °C for 15 min, hold at 4 °C until further use.

Next, 2 uL T4 DNA Ligase (Enzymatics, Catalog #L6030-LC-L) and 1 uL of one of the 10 uM annealed Y adapters (chosen from a 96-well plate of GS_i5 adapters) was added per end-repaired and A-tailed reaction. Each reaction was mixed and incubated on a thermocycler with the following program: 16 °C for 30 min, 22 °C for 30 min, hold at 4 °C until further use.DNA was then purified using 0.9 volumes Ampure XP beads using the same bead clean-up protocol but using 23 uL IDTE pH 7.5 to resuspend the DNA-bead mixture for elution and collecting 22 uL supernatant after incubation on the magnet.

Next, ligated and sheared DNA fragments were amplified using a primer specific to all adapters (P5_1) and a primer specific to the sequence of interest (GSP1 +/−). To each tube on ice, the following reagents were added: 22 uL DNA from previous step, 3 uL 10X Platinum Taq Polymerase PCR Rxn Buffer (-Mg2 free) (Invitrogen, Catalog #10966034), 1.2 uL 50 mM MgCl2 (Invitrogen, Catalog# 10966034), 0.6 uL 10 mM dNTP mix (Invitrogen, Catalog #18427013), 0.5 uL 10 µM P5_1 primer, 1 uL 10 µM PSP1+/−, 1.5 uL 0.5 M TMAC (Sigma Aldrich, Catalog #T3411), and 0.3 uL Platinum Taq DNA polymerase (5 U/μL) (Invitrogen, Catalog #10966034) to a total of 30.1 uL. Each reaction was mixed and incubated with the following thermocycler conditions: initial melt of 95 °C for 2 min; 15 cycles of 95 °C for 30 s, 70 °C (−1 °C/cycle) for 2 min, and 72 °C for 30 s; followed my 10 cycles of 95 °C for 30 s, 55 °C for 1 min, and 72 °C for 30 s; followed by a final extension at 72 °C for 5 min; hold at 4 °C until further use.

Amplified DNA was purified using the bead clean-up protocol but with 1.2 volumes of Ampure XP beads and using 21 uL IDTE pH 7.5 to resuspend the DNA-bead mixture for elution and collecting 20.4 uL supernatant after incubation on the magnet.

Next, the second round of PCR amplification was performed to add plate barcodes (GS_i7 sequences). To each tube on ice, the following reagents were added: 1.5 ul 10 µM Plate adapter (GS_i7), 20.4 ul DNA from previous step, 3 ul 10X Platinum Taq Polymerase PCR Rxn Buffer (-Mg2 free) (Invitrogen, Catalog #10966034), 1.2 ul 50 mM MgCl2 (Invitrogen, Catalog #10966034), 0.6 ul 10 mM dNTP mix (Invitrogen, Catalog #18427013), 0.5 ul 10 µM P5_2 primer, 1 ul 10 µM PSP2+/−, 1.5 ul 0.5 M TMAC (Sigma Aldrich, Catalog #T3411), and 0.3 ul Platinum Taq DNA polymerase (5 U/μL) (Invitrogen, Catalog #10966034) to a total of 30 ul. Each reaction was mixed and incubated with the following thermocycler conditions: initial melt of 95 °C for 5 min; 15 cycles of 95 °C for 30 s, 70 °C (−1 °C/cycle) for 2 min, and 72 °C for 30 s; followed by 10 cycles of 95 °C for 30 s, 55 °C for 1 min, and 72 °C for 30 s; followed by a final extension at 72 °C for 5 min; hold at 4 °C until further use.

Afterwards, 25 uL of each barcoded reaction was combined into one pool and 0.7 volumes of Ampure XP beads were added. Samples were mixed 10 times and incubated for 5 minutes at room temperature. They were then added to the magnet for 5 minutes then the supernatant was discarded. 2 1x volumes of freshly made 70% ethanol were added, incubating for 30 seconds each time before discarding the supernatant. After the last wash, the beads were air-dried for 6-8 minutes. 75 uL IDTE pH 7.5 was added and the tubes were removed from the magnet and mixed, then incubated for 2 minutes. The reaction was separated on the magnet for 2-4 minutes and then the supernatant was collected into a new tube for NGS library sample submission. Pooled eluate was quantified using the Qubit and the Qubit dsDNA HS kit following the recommended protocol.

Final products were sequenced on a MiSeq or NextSeq 2000 with paired-end 150 bp reads with the cycle settings: 148-10-22-148 for MiSeq or 151-10-22-151 for NextSeq. Samples were sequenced to obtain at least 3,000,000-fold coverage per sample. For MiSeq reactions, 3 uL of 100 uM custom sequencing primer Index1 (5’-3’: ATCACCGACTGCCCATAGAGAGGACTCCAGTCAC) was added to MiSeq Reagent cartridge position 13 and 3 uL of 100 uM custom sequencing primer Read2 (5’-3’: GTGACTGGAGTCCTCTCTATGGGCAGTCGGTGAT) was added to MiSeq Reagent cartridge position 14.

For NextSeq reactions, 1.98 ul of 100 uM custom sequencing primer Read2 was added to 600 ul Illumina HP21 primer mix for a 0.3 uM final concentration and 3.98 ul of 100 uM custom sequencing primer Index1 was added to 600 ul BP14 primer mix for a 0.6 uM final concentration. Then 550 μl of custom primer mix was added to custom 1 well or custom 2 well on the reagent cartridge separately.

### T cell Transfection

Primary human T cells were thawed on day -3 and activated with anti CD3/CD28 beads (Thermo Fisher Scientific, Catalog #CTS40203D). Three days later, ‘day 0’, T cells were centrifuged at ∼300 x g for 10 min, and washed with 1x DPBS (Corning, Catalog #21-031-CV). For 96-well nucleofection, cells were resuspended at 1E6 cells per 20 µl of P3 Primary Cell Nucleofector Solution (Lonza, Catalog #V4SP-3096) with supplement. 20 µl of cells were mixed with 9.6 µl of Bxb1 mRNA and nanoplasmid donor (Aldevron), then transferred to the Lonza 96-well Shuttle plate. Nucleofector program EO-115-AA was used to electroporate T cells with the mRNA/DNA mix on the Amaxa Nucleofector 96-well Shuttle System (Lonza). After electroporation, cells were incubated for 10 min at room temperature, 80ul of T-cell medium were added per well and cells transferred to a 48-well tissue culture plate containing 400 µl medium (prewarmed to 37 °C). Electroporated T cells were placed in a 30°C incubator for 24 hours and then transferred to a 37°C incubator. Cells were cultured until day 14 by replenishing culture medium every 3 to 4 days. On day 14, GFP expression was determined by Flow Cytometry and T cell aliquots were harvested, DNA isolated and PCR amplified with the specified primers to assess target integration into the TRAC by Illumina next generation sequencing.

### Flow Cytometry of T cells

On day 14 post transfection, ∼5×10^5 cells were harvested, washed once with PBS, then stained with Fixable Viability Dye eFluor™ 780 (Thermo Fisher, Catalog #65-0865) according to Manufacturer’s instructions. The GFP flow analysis was performed using an Attune NxT Acoustic Cytometer (Thermo Fisher Scientific). Subsequently, the flow data was analyzed with FlowJo software (BD).

For cell sorting 3ml (∼ 6E6) cells were spun down, washed once with PBS, de-beaded using a Dynal MPC-L magnetic particle concentrator (Thermo Fisher) and then resuspend in 1.5ml of EasySep buffer (Stemcell Technologies, Catalog #20144). The cell suspension was loaded on a Sony cell sorter SH800S for sorting & GFP fraction collection. For the sorting, a 70um sorting chip (Sony, Catalog #LE-C3207) and sorting pressure 4 was employed. A small volume of the mock transfected control sample was used to set the GFP negative gate and a MINT transfected sample to set gates for the different expression levels of GFP as shown in Extended Data Fig. 11.+. For cell collection, 0.4 ml PBS were pre-aliquoted to each 15 ml collection tube, desired numbers (when available) of GFP negative & GFP positive fractions were collected.

After GFP fractions were collected, an aliquot of the sorted samples was centrifuged and appropriate GFP fractionation was confirmed by flow cytometry analysis. The remaining cells were lysed with QuickExtract buffer for DNA extraction and analyzed for TI as described above.

### Computational analysis of directed evolution experiments

Data processing was performed with custom python scripts and logo plots were generated using Logomaker^62^. Sequence reads that don’t match sequences encoded by the directed evolution library were filtered out as were sequence reads that were likely artifacts due to being rare sequence reads with only one or two differences from much more common sequence reads. The remaining sequence reads were translated into peptide sequence, and the rest of the analysis only considered the peptide sequence of the randomized region. For hairpin selections, the partially randomized position 322 could be either a G, A, R, or P and the motif analysis was conducted separately based on the identity of position 322. The clearest signal was 4 residue motifs within the 6 fully randomized positions. For example, the helix library with a XXXX<u>L</u>XX randomization scheme had a strongly enriched motif of XGGN<u>L</u>XR where X is any amino acid. To account for potential biases in the starting library, motifs were scored based on the FDR corrected p-value of a given motif occurring by chance given the amino acid frequency at each position when considered independently. For helix selections, enriched 4-residue motifs that were observed in selections with a wide variety of different DNA target sequences were considered to be non-specific and were filtered out. Compatible four residue motifs (e.g. XGXN<u>L</u>KR and XXGN<u>L</u>KR) were then combined to create motifs. Several peptides that best matched the most enriched 4 residue motifs in each combined motif were then chosen for additional characterization.

### Computational analysis of pooled screening of attB sites in K562 cells

A custom python script was used to count sequence tags corresponding to different recombined targets. Counts were normalized to the total number of sequence reads obtained for a given sample. Normalized sequence tag counts for a given Bxb1 variant + wild-type Bxb1 at a given site were then compared to the normalized sequence tag counts for wild-type Bxb1 alone at for the same sequence tag.

### Computational analysis of chromosomal targeted integration events

Sequence data was first processed using our indel analysis software pipeline. The output from this analysis was further processed using a custom python script that identified aligned sequence reads that contained the sequence from the right attP half-site from the donor. Sequence reads that contained this integration tag, but were not the expected length were scored as “TI + indel”. Sequence reads that contained this integration tag and were the expected length were scored as “perfect TI”. Sequence reads that did not contain the integration sequence tag were scored as either wild-type amplicon or non-TI indel based on the output of the indel analysis software.

### Computational identification of Bxb1 pseudo-sites

Raw data from Bessen et al. was processed to produce a position weight matrix for the Bxb1 attB and attP target sites. A custom python script scanned the human genome for potential target sites that matched the strongly preferred G nucleotide at position −4 and at position +4. Sequences that met these criteria were then scored against the position weight matrix. The left and right half-sites of the natural Bxb1 attB site are quite different from each other and likely represent different binding modes of the Bxb1 hairpin region. Thus, for attB sites, each potential site was scored against a position weight matrix representing an inverted repeat of the left half-site of the natural attB sequence, and inverted repeat of the right half-site of the attB sequence, the composite attB left and right half-sites on the top strand of DNA, or the composite attB left and right half-sites on the bottom strand of DNA. The top 12 scores for each category were experimentally characterized for a total of 48 potential attB pseudo-sites in the human genome. The natural attB site is much more symmetric so the position weight matrix for the left and right half-sites was averaged together and this averaged position weight matrix was used to score both sites of potential attP pseudo-sites in the human genome. The top 48 scoring potential attP pseudo-sites were also characterized experimentally.

### Computational analysis of the unbiased genome-wide specificity assay

NGS reads were demultiplexed, adapter trimmed, and filtered for a minimum quality threshold of 14 over all bases. Samples then underwent analysis for plasmid integration site detection. NGS samples were processed to remove remaining contaminant unintegrated plasmid reads due to incomplete DpnI digestion or no DpnI digestion in the case of week 3 and 4 samples. Reads were then aligned to the hg38 genome, and candidate integration sites were summarized. First, read 2 reads that contained both attP sequence 5’ and 3’ of the dinucleotide were removed from analysis, corresponding to unintegrated donor plasmid reads. Then all sequence up to the start of the dinucleotide (up to and including the 5’ attP sequence) was removed, leaving the remaining sequence to align to the hg38 genome using Bowtie2. Alignments with a MAPQ less than 23 were removed from the analysis. Next, common read1 start locations, which correspond to unique genomic shear locations and ligation events generated in the protocol, were used to deduplicate common reads. Unique read1 start and dinucleotide positions were then summed per dinucleotide position to generate a list of deduplicated reads per potential integration event. Next, all reads per reaction were summed per alignment position in the genome and per alignment orientation (top and bottom strands). Then, positions were combined into a single potential integration location per reaction and alignment orientation if they fell within a 50 bp window of one another, all reads per this grouping were summed and the coordinate with the most reads was kept per group. Lastly, potential integration locations across top and bottom strand alignments and between both “plus” and “minus” reactions per original transfected sample were combined. Common potential integration locations were merged into one potential integration location if within 50 bp of one another (this would encompass alignments separated by a dinucleotide as a result of sequencing upstream and downstream of integration in both reactions). The final list of potential integration loci was inspected for expected integration genotypes (2 merged locations, in opposite alignment orientation, separated by a dinucleotide that corresponds to the donor plasmid dinucleotide used in the assay).

### Single cell cloning and targeted integration characterization

K562 cells were nucleofected with eebxb1-ZFP constructs as detailed above. Three days post-nucleofection, cells were examined via microscopy and quantified using automated cell counters (Thermo Fisher Scientific). Subsequently, cells were seeded into 96-well plates at an average density of 0.3 cells per well. On day 10, wells exhibiting GFP fluorescence and containing a single cell cluster were maintained for an additional 10 days. Thereafter, cells were subjected to flow cytometry analysis and genomic DNA was extracted using QuickExtract (Lucigen #QE09050) for TI analysis.

To verify full-length and single-copy donor integration, DNA from single-cell clones demonstrating >99% target integration efficiency and 100% GFP positive was amplified using primers LL-P125 (5’-GATCAATCTGAGTGCTGTACGG-3’) and LL-P126 (5’-ACCCCTGTCTTACCTGTTTCAA-3’). Briefly, PCR reactions were carried out in 20uL volumes, using 2 x Phusion High-Fidelity PCR master mix (NEB M0531L), and 2 uL of quick extracted DNA as template. Thermocycling was carried out as follows 98 °C 1min; 35 cycles of [98 °C 15sec, 65 °C 30sec, 72 °C 10min] Final extension 72 °C 5 min. The resulting PCR products were resolved by gel electrophoresis and confirmed by Nextera sequencing following the instruction of Illumina Nextera XT DNA Library Preparation Kit (FC-131-1096).

---

**Extended Data Figure 1.**
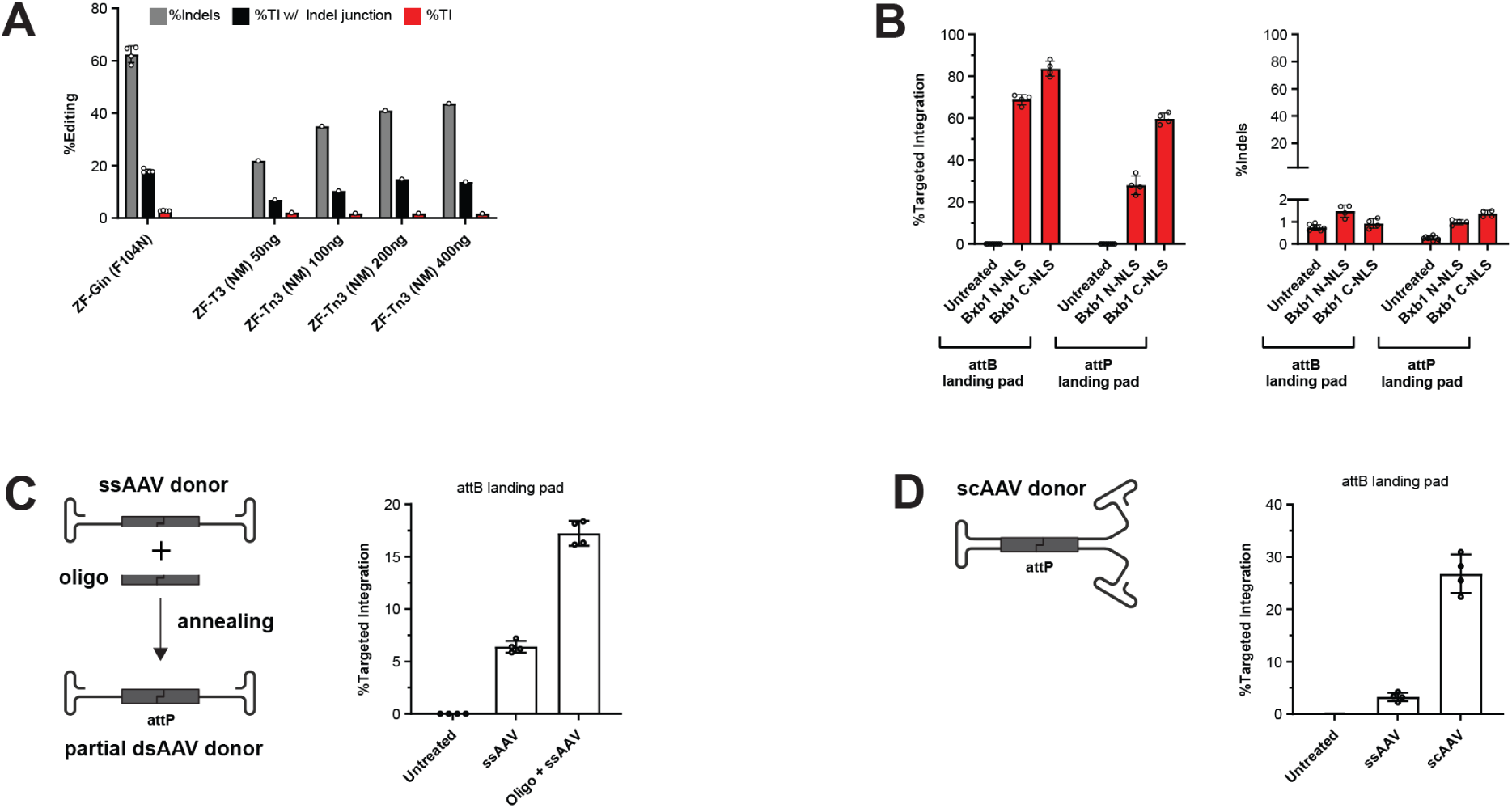
Performance of zinc fingers fused to small serine recombinase domains and performance of different wild-type Bxb1 constructs and donors. **a.** Performance of Zinc Finger (ZF)-targeted small serine recombinase Gin and Tn3 hyperactive domains. The data shown in this graph is derived from separate experiments. In contrast to Bxb1, both Gin and Tn3 hyperactive domain fusion proteins can result in high levels of indels causing low product purity and hindering further improvement of target integration frequencies. We also observed indels within the assayed TI junction site. Data is derived from a PCR-based NGS. Data are presented as the mean from four biological replicates for ZF-Gin, and individual data points are shown for ZF-Tn3 fusions. **b.** Performance of wild-type Bxb1 against natural attB and attP target sequences in human cells. We first established K562 landing pad cell lines by installing the natural attB or attP sequence in the human AAVS1 locus. We noticed improved performance of Bxb1 with a C-terminal NLS compared to the construct with a N-terminal NLS. This guided future Bxb1 designs where all evolved variants presented in this study have a C-terminal NLS. We also noticed higher TI into the attB landing pad. Notably, no or only minimal levels of indels were observed within the landing pad target sequences. Data is derived from a PCR-based NGS. Data are presented as the mean ± s.d. from four biological replicates (eight for untreated sample). **c.** Utilization of a single-stranded AAV (ssAAV) donor for Bxb1-mediated TI. We tested ssAAV as a donor against an attB landing pad in K562 cells and noticed measurable integration levels that can be increased through co-delivery of an oligonucleotide that is complementary to the attP sequence, therefore making the ssAAV donor partially double-stranded. **d.** Utilization of a self-complementary AAV (scAAV) donor for Bxb1-mediated TI. Data are presented as the mean ± s.d. from at least four biological replicates.

**Extended Data Figure 2.**
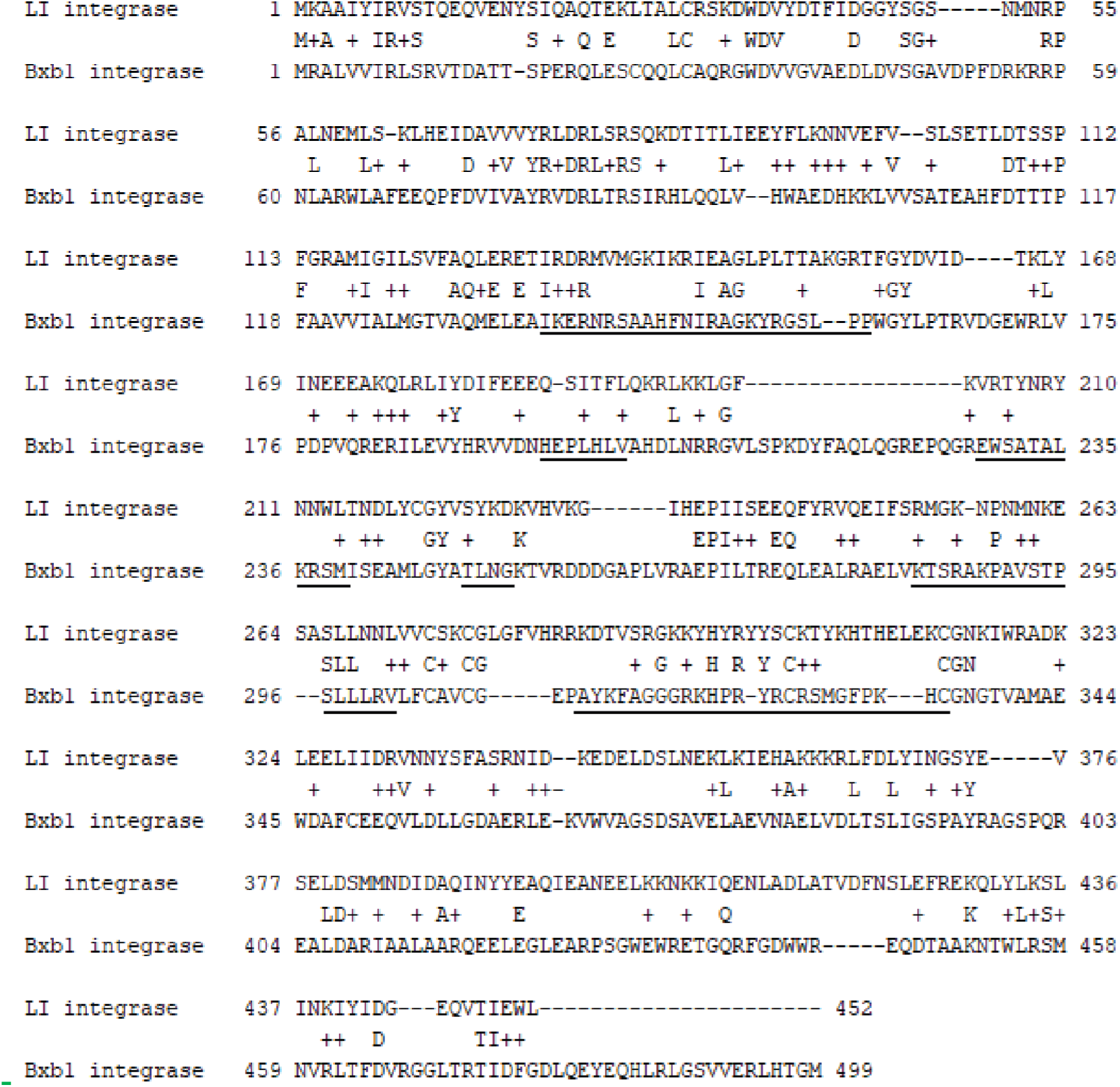
Sequence alignment of the large serine recombinase from the Listeria innocua prophage and Bxb1. The alignment was modified to reflect predicted secondary structures. The region of Bxb1 that was probed with saturation mutagenesis is underlined.

**Extended Data Figure 3.**
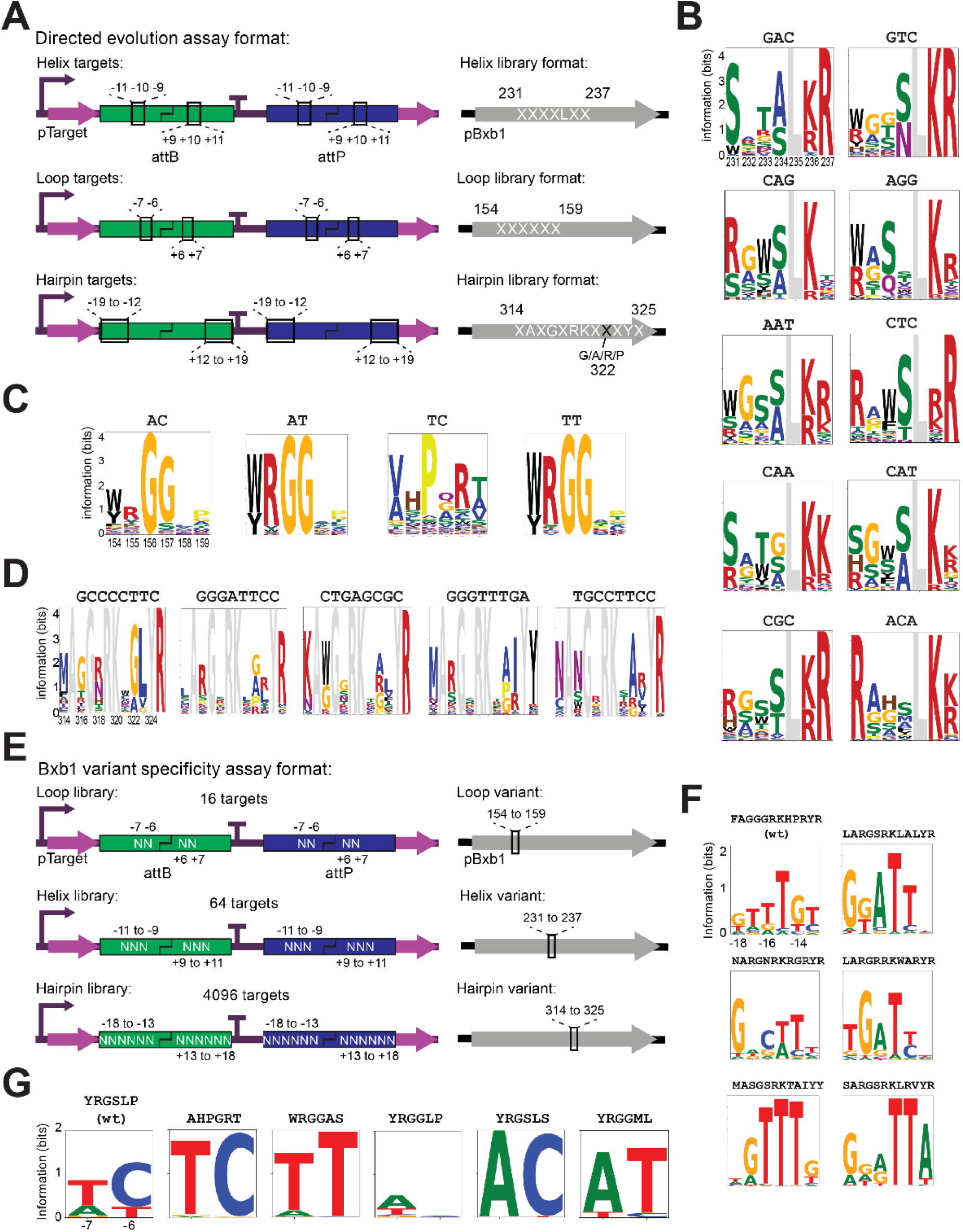
**a.** Integrase variant library screening configuration with pBxb1 libraries where the helix, loop, or hairpin submotif has been randomized and are transformed along with pTarget plasmids where corresponding positions of the attB and attP target sites are altered. Primers with NNK codon usage were used for randomization of each residue indicated with an X to generate the Bxb1 variant libraries. For the hairpin library, SSK was used instead at residue 322 to bias the randomization to an A or P at this position. **b.** Example sequence logo plots summarizing enriched peptide motifs at amino acid residue positions 231-237 for helix selections with the corresponding helix DNA targets for each selection shown above each sequence logo. Position 235 was not randomized during selection and is shown in grey. Helices with an arginine (R) at position 237 were frequently observed in selections for a C at position −9 of the DNA target, helices and a lysine (K) at position 237 were often observed in selections for a A at position −9 of the DNA target, helices with an asparagine (N) at position 234 were often observed in selections for a T at position −10 of the DNA target, helices with an alanine (A) or glycine (G) at position 234 were often observed in selections for A at position −10 of the DNA target, and a tryptophan (W) at position 233 was often observed in selections with a C at position −11 of the DNA target. These correlations mimic interactions observed with engineered zinc fingers if one considers position 233 to correspond to +2 of the zinc finger recognition helix, 234 to correspond to +3 of the zinc finger recognition helix, and 237 to correspond to +6 of the zinc finger recognition helix and if one considers the zinc finger DNA triplet to be the reverse complement of the Bxb1 helix target triplet. For example, from Ichikawa *et al.* (2023), the zinc finger helix QSGTLR**<u>R</u>** can target **<u>G</u>**CA, TKAYLL**<u>K</u>** can target **<u>T</u>**GA, QSS**<u>N</u>**LRT can target A**<u>A</u>**A, DPS**<u>A</u>**LIR can target A**<u>T</u>**C, and RK**<u>W</u>**TLQQ can target AA**<u>G</u>.** This implies some similarities between how the Bxb1 helix and how the zinc finger recognition helix interacts with target DNA. **c.** Example logo plots summarizing the results of selections at amino acid positions 154-159 for each loop DNA target selection shown above each sequence logo. **d.** Example logo plots summarizing the results of selections at amino acid positions 314-325 for each hairpin DNA target selection shown above each sequence logo. Residues that were not randomized are shown in gray. **e.** Bxb1 variant specificity assay configuration where a single integrase variant previously isolated in the directed evolution assay is assayed for activity across DNA target libraries varied at the indicated positions with N. Each individual pTarget in the library contains the same modification at both half-sites of the attB and attP sites. **f.** Example DNA specificity plots of selected hairpin variants. Shown is the mean of a total of eight replicates; four biological replicates and two technical replicates thereof. **g.** Example DNA specificity plots of selected loop variants. Shown is the mean of four biological replicates.

**Extended Data Figure 4.**
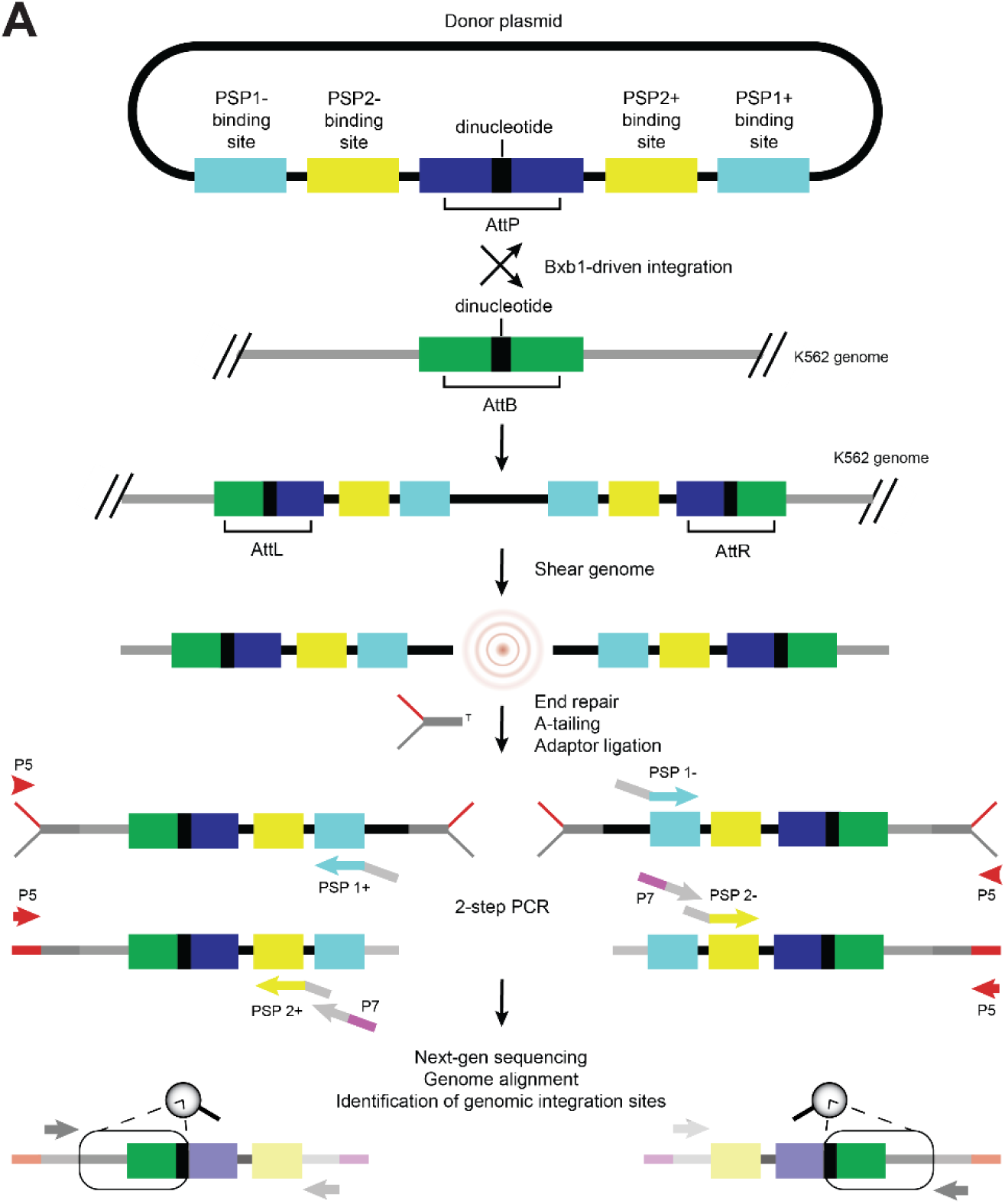
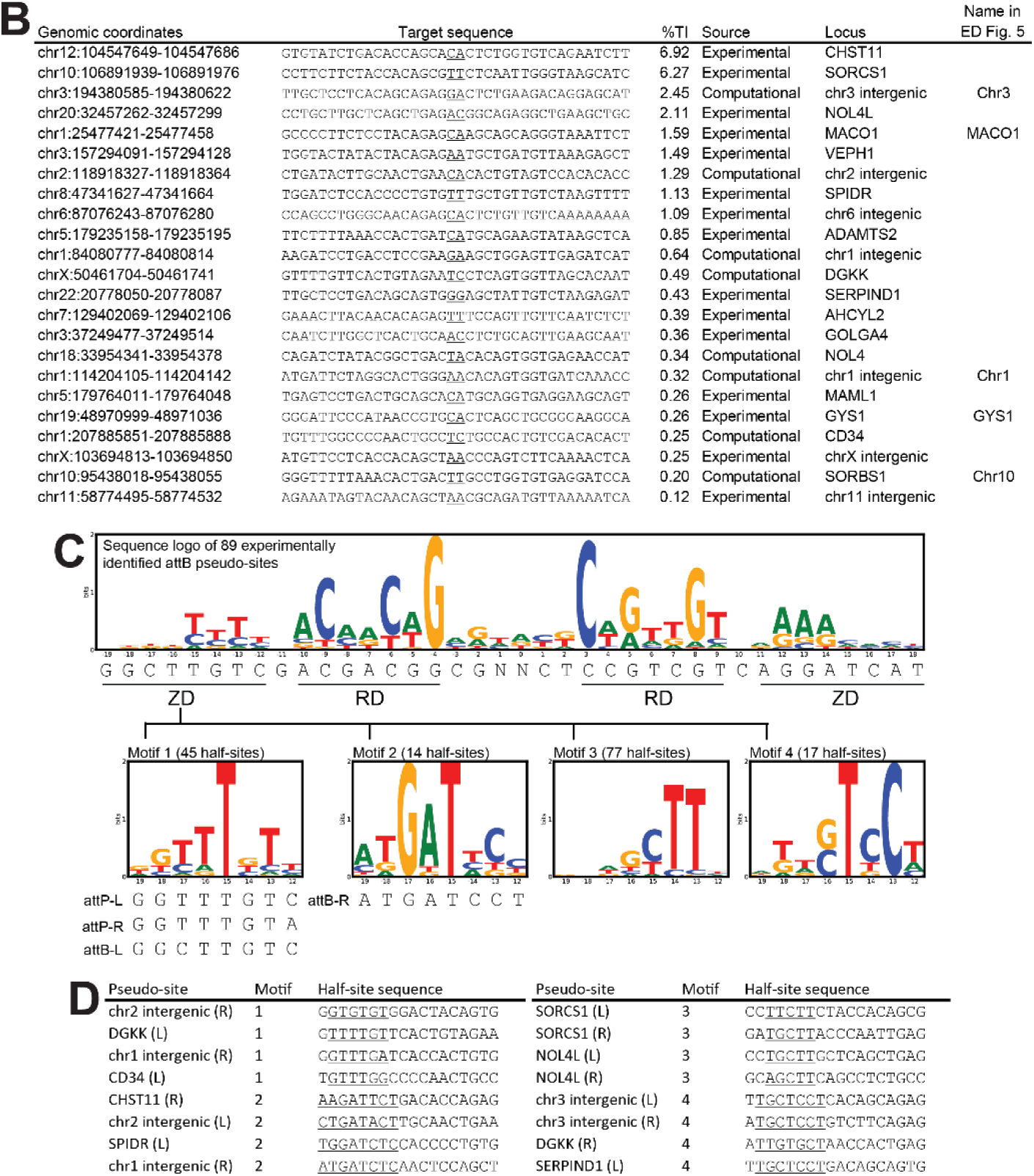
Identification of Bxb1 pseudo-sites in the human genome using an unbiased genome-wide specificity assay. **a.** Schematic of the unbiased genome-wide specificity assay used in this study to experimentally identify potential Bxb1 integration sites in the human genome. Bxb1 drives integration of a donor plasmid at the intended on-target and pseudo-sites throughout the K562 genome. DNA is then extracted, sheared and adapters are ligated. Both genome-plasmid junctions are amplified separately using a 2-step PCR with plasmid-specific primers (PSP1 and 2) from either upstream or downstream of the original AttP site (+ or -). Then genomic integration sites are identified using NGS and a custom bioinformatics pipeline. **b.** List of 23 validated Bxb1 pseudo-sites in the human genome that were either identified through a computational search or experimentally using the assay shown in panel **a**. The central dinucleotides in each site are underlined. Only the portion corresponding to an attB site are shown in the table. Site-specific donors were used to validate TI using a PCR-based NGS assay. CD34 and Chr1 pseudo-sites were identified in an independent computational search designed to identify targets for Bxb1 variants; control experiments indicated that these sites were better targeted with wild-type Bxb1. **c.** Sequence logo for all 89 potential pseudo-sites with at least two independent integrations identified using the experimental assay shown in panel **a**. Positions −19 to −12 of the 178 half-sites of these 89 potential pseudo-sites are a mixture of four different distinct DNA motifs and sequence logos for these four different motifs are shown below the sequence logo of the full sites. **d.** Examples of half-sites from validated human pseudo-sites for wild-type Bxb1 that correspond to each of the four DNA sequence motifs shown in panel **c**. Positions in each half site that correspond to key bases in the relevant motif are underlined.

**Extended Data Figure 5.**
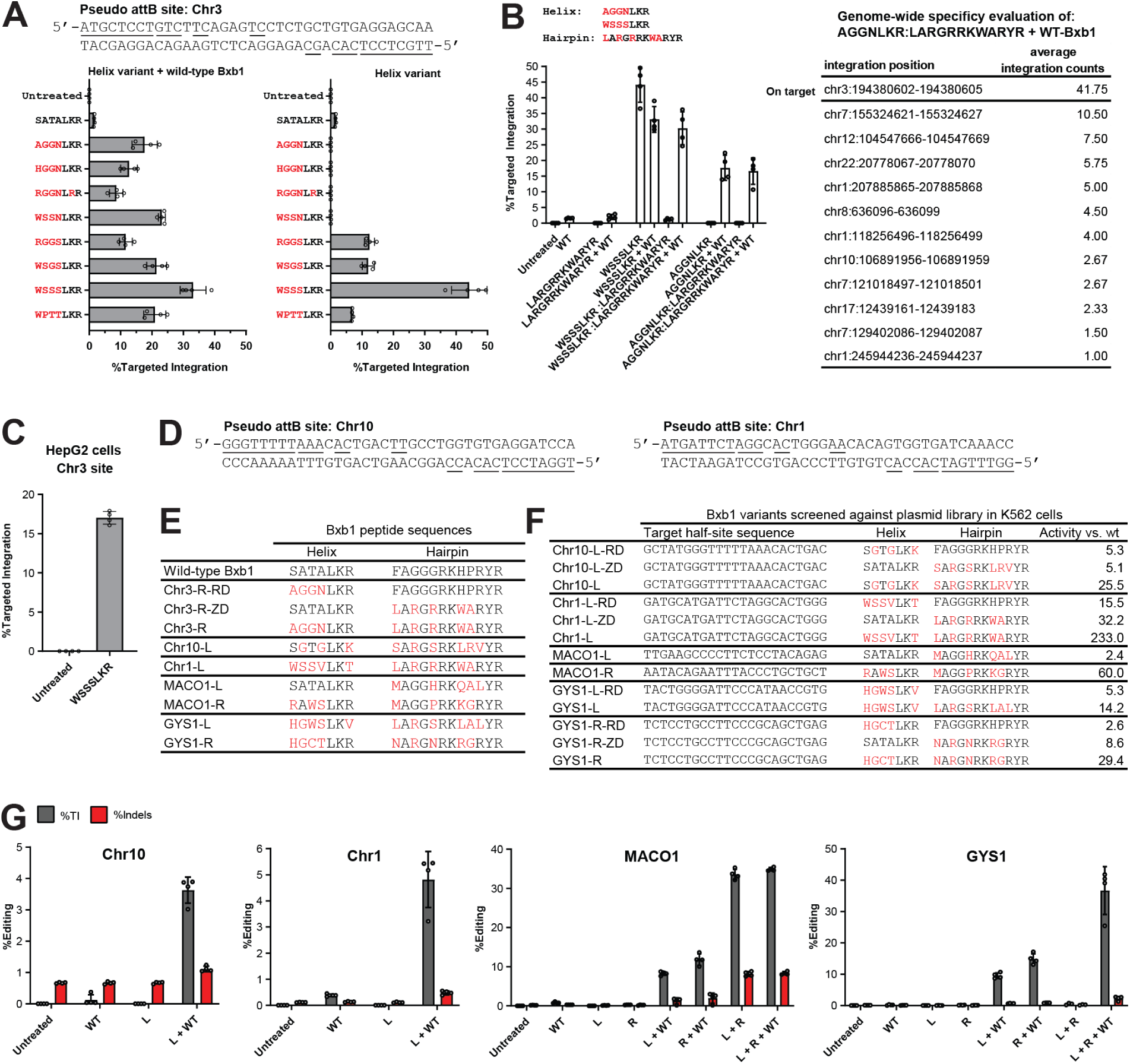
Performance of Bxb1 variants against pseudo-sites in the human genome. **a.** The attB pseudo-site on chromosome 3 used for initial targeting and performance measurements of various Bxb1 helix domain variants against this target site using a PCR-based NGS assay. SATALKR is the wild-type helix peptide sequence and the target DNA sequence is shown as the reverse complement of the sequence in Extended Data Fig. 4b to make it easier to visualize the region we are targeting in this experiment. Selected helix sequences that match enriched motifs and contain an S or T at position 234 were active at the chromosome 3 target even in the absence of wild-type Bxb1 indicating that such helices can interact with both half-sites in the genomic target as well as the attP target in the donor. Similar sequences were enriched in selections with other target sites, so we concluded that these types of helix sequences represent helices with poor specificity, and we computationally removed similar sequences from subsequent selections. In contrast, selected Bxb1 variants with an N at position 234 such as AGGNLKR were only active at the chromosome 3 target when mixed with wild-type Bxb1. DNA targeting specificity characterization of this helix indicated a dramatic shift in specificity towards a T at position −10 (Figure 1f) consistent with the GTC sequence at positions −11 to −9 of the half-site we targeted with this helix selection **b.** On-target performance of a Bxb1 variant with combined helix and hairpin variations, and data summary of a genome-wide specificity evaluation using a modified version of the assay described in Extended Data Fig. 4 (see Extended Data Fig. 6). Here, we combined the most promising selected hairpin LARGRRKWARYR with both the AGGNLKR helix and the more active, but less specific WSSSLKR helix. Both combinations retained full activity at the chromosome 3 site when combined with wild-type Bxb1 but show a substantial decrease without wild-type Bxb1. Thus, WSSSLKR in combination with LARGRRKWARYR now requires wild-type Bxb1 to achieve full integration activity. This indicates improved specificity vs. the WSSSLKR helix variant alone which is able to interact with both half sites of the chromosome 3 target as well as the attP site in the donor in the absence of wild-type Bxb1. Since both AGGNLKR and LARGRRKWARYR were able to target the intended chromosome 3 half-site more specifically than the wild-type Bxb1 helix and hairpin, we reasoned the combination of this helix and hairpin should be even more specific for the chromosome 3 site and characterized this combination of helix and hairpin using an unbiased genome-wide specificity assay. This fully engineered Bxb1 variant had fewer candidate integration sites within the human genome than wild-type Bxb1 and we observed that the intended site on chromosome 3 had higher levels of integrations than any other site in the human genome. Note that the indicated Bxb1 variant has to be mixed with wild-type Bxb1 (included to bind the attP site on the donor) in order to be active at the intended chr3 target site so the genome-wide specificity assay was performed with a mixture of the indicated Bxb1 variant and wild-type Bxb1. The experiment was performed with a pool of 16 donor constructs containing all possible central dinucleotides. Average integration counts were determined using the average deduplicated read numbers across 2 samples using primers that amplify the attL and attR plasmid-genome junctions. The corresponding control sample where GFP was substituted for Bxb1 resulted in 0 sites (Supplementary Table 21) **c.** Activity-testing of a Bxb1 helix variant from panel a. in human HepG2 cells. **d.** Sequence of two additional Bxb1 pseudo-sites in the human genome. **e.** Bxb1 peptide sequences of evolved Bxb1 variants that showed improved performance against the half-sites of pseudo-sites shown in Figure 2b and panel c. **f.** Screening data using synthetic DNA targets tested in human K562 cells that was used to identify the constructs shown in panel d. Activity is determined by the number of DNA sequence reads corresponding to recombined versions of each synthetic target; activity is normalized to the activity of wild-type Bxb1 alone against the same synthetic target site. **g.** Results from a PCR-based NGS assay demonstrating improved performance of evolved Bxb1 variants against their chromosomal endogenous targets in K562 cells. The presence of a wild-type Bxb1 expression construct is necessary to bind the wild-type attP sequence on the donor plasmid. Data are presented as the mean ± s.d. from four biological replicates.

**Extended Data Figure 6.**
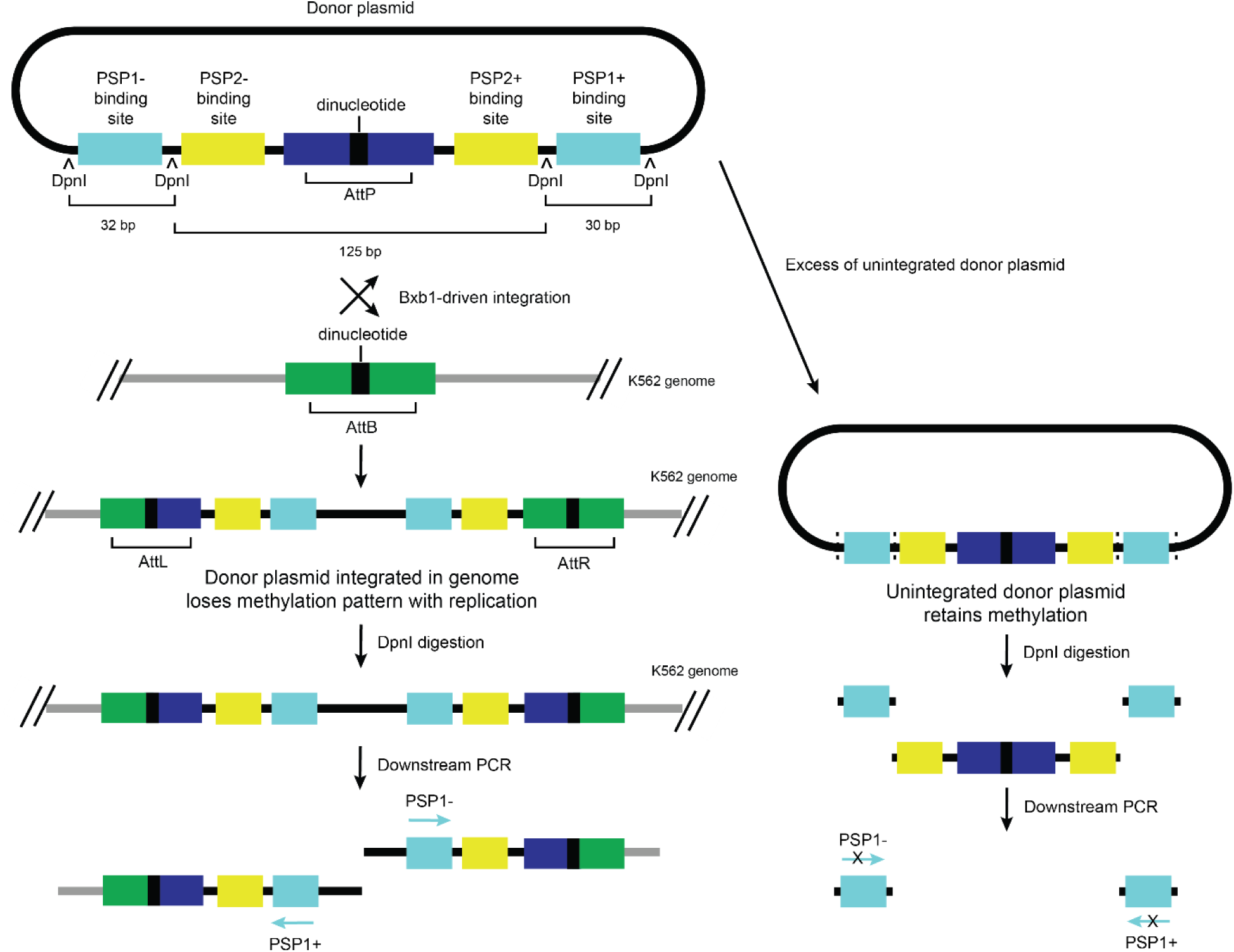
Improved genome-wide specificity assay. Schematic of the modified unbiased genome-wide specificity assay used in this study to experimentally nominate candidate integration sites shown in Extended Data Fig. 5b. Strategically placed DpnI recognition sites in the donor molecule supports the enzymatic removal of excess unintegrated donor plasmid since these sites should lose their bacterial methylation marks after integration and cell replication in the human genome. The DpnI removal method resulted in a substantial reduction of donor plasmid-derived background signal and enabled culturing the cells for less time compared to previous method where many cell passages were required to dilute out excess plasmid.

**Extended Data Figure 7.**
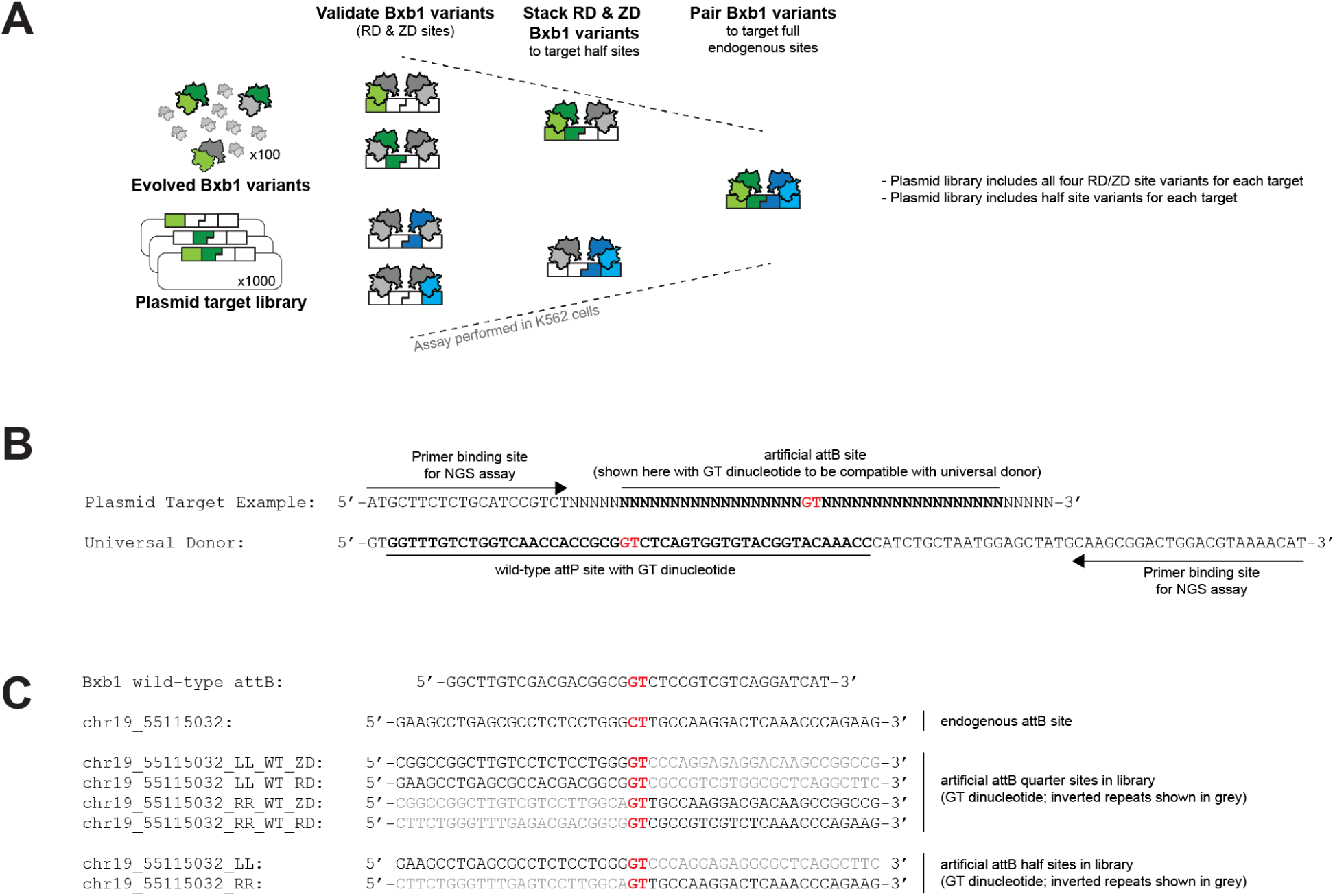
Screening system for testing Bxb1 variants against a plasmid library of artificial target sites in human K562 cells. **a.** Schematic of a system for testing evolved Bxb1 variants against thousands of artificial targets in a plasmid target library as described in Figure 3a**. b.** Sequence examples for the target library and the donor, including assay design information for PCR-based detection of recombination events. **c.** Sequence example for all plasmid target library members of the AAVS1 site highlighted in Figure 3. The plasmid target library includes up to six distinct members for each endogenous target site. Four members are designed to support the identification of Bxb1 variants targeting the RD and ZD motifs of both endogenous half-sites. Two additional members are designed to confirm activity of stacked Bxb1 variants against the corresponding half-sites. Bxb1 variants are screened individually against the plasmid target library.

**Extended Data Figure 8.**
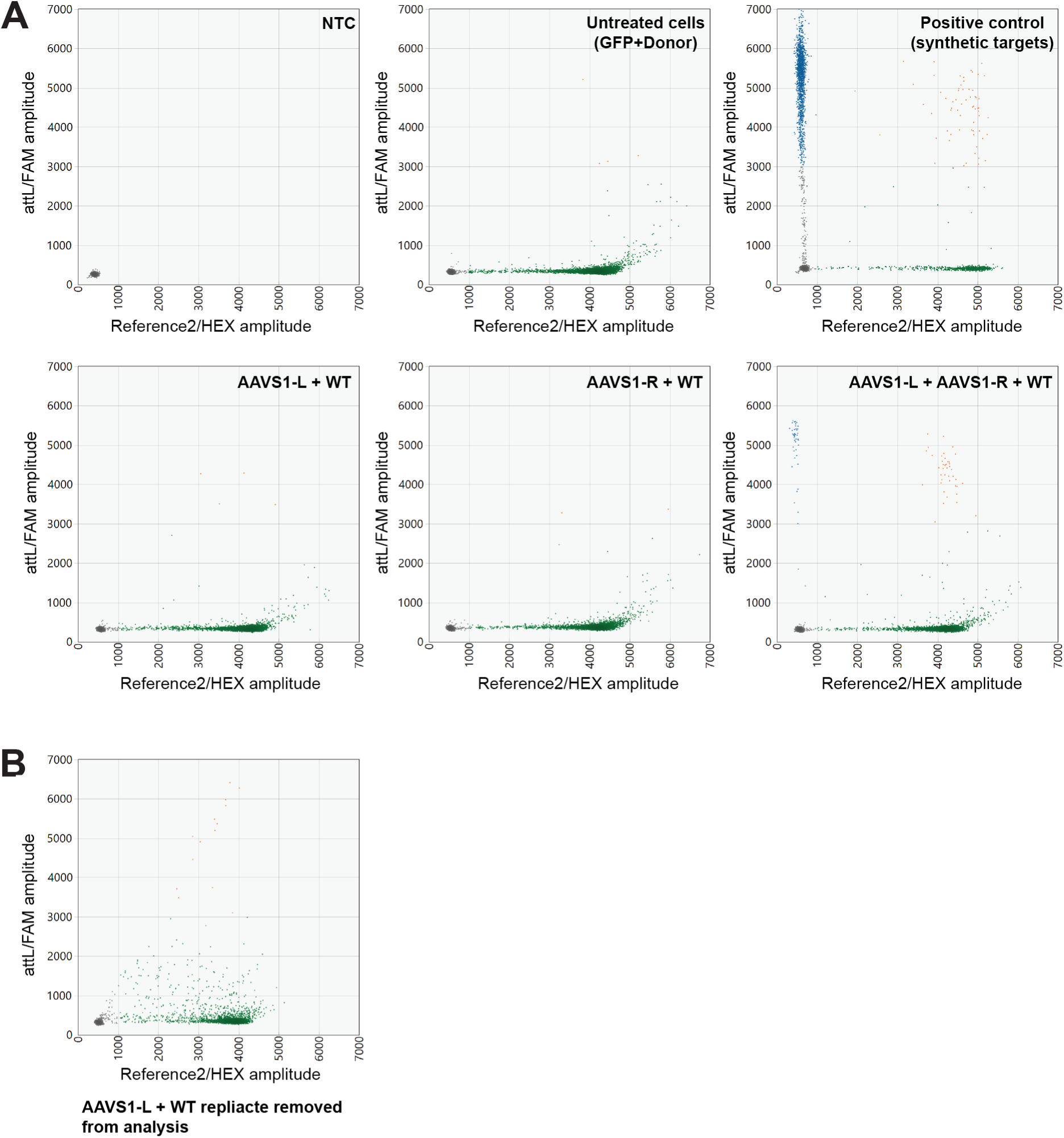
ddPCR analysis of Bxb1 variants targeted to the human AAVS1 locus. **a.** Example 2-D ddPCR scatterplots of differently treated samples. Each 2-D plot contains 1-4 clusters of droplets: (1) double-negative droplets containing no targeted DNA templates (grey dots clustered at the left bottom in each plot); (2) reference-only droplets (green); (3) target integration/attL-only droplets (blue); and (4) double-positive droplets containing both target integration/attL and reference DNA templates (orange). **b.** The 2-D plot of an AAVS1-L + WT replicate discarded from the analysis with noisy FAM signal likely due to shredded droplets.

**Extended Data Figure 9.**
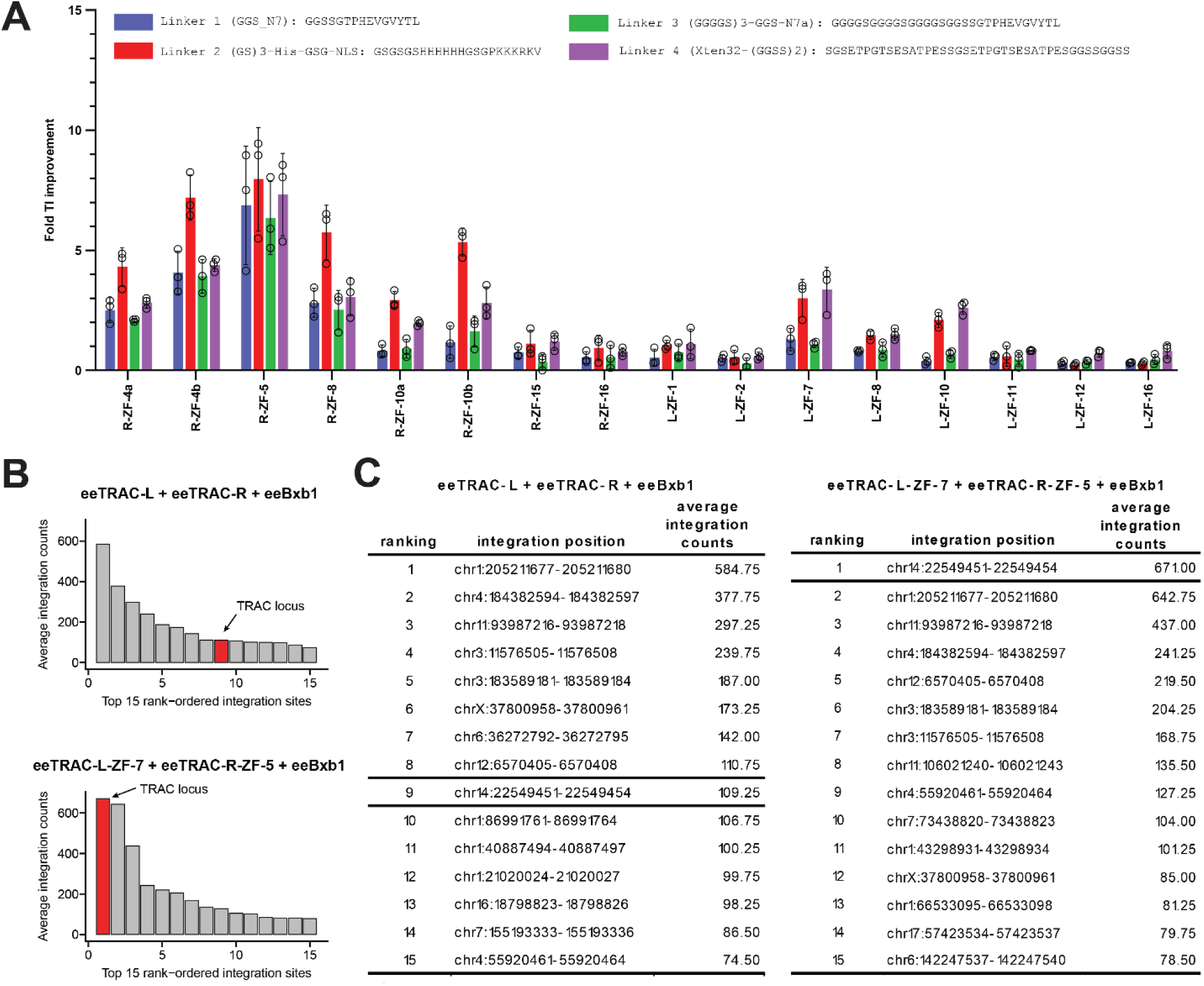
ZF and activity-increasing TRAC Bxb1 construct optimizations. **a.** Four linkers with different lengths were tested in the context of the constructs highlighted in Figure 4. Linker 2 was chosen for all subsequent experiments. **b.** Results from an unbiased genome-wide specificity analysis of the indicated constructs that contain the eeBxb1 mutations. The top 15 candidate integration sites are shown (out of total 2637 sites for eeTRAC constructs and 1949 for ZF-fused eeTRAC constructs) with the on-target at the TRAC locus indicated between bolded lines. The entire datasets are provided in Supplementary Tables 18 and 19 and corresponding control datasets in Supplementary Tables 17 and 20. Average integration counts were determined using the average deduplicated read numbers across 2 replicates and 2 samples each using primers that amplify the attL and attR plasmid-genome junctions. **c.** Corresponding table format of data in b with candidate integration site information.

**Extended Data Figure 10.**
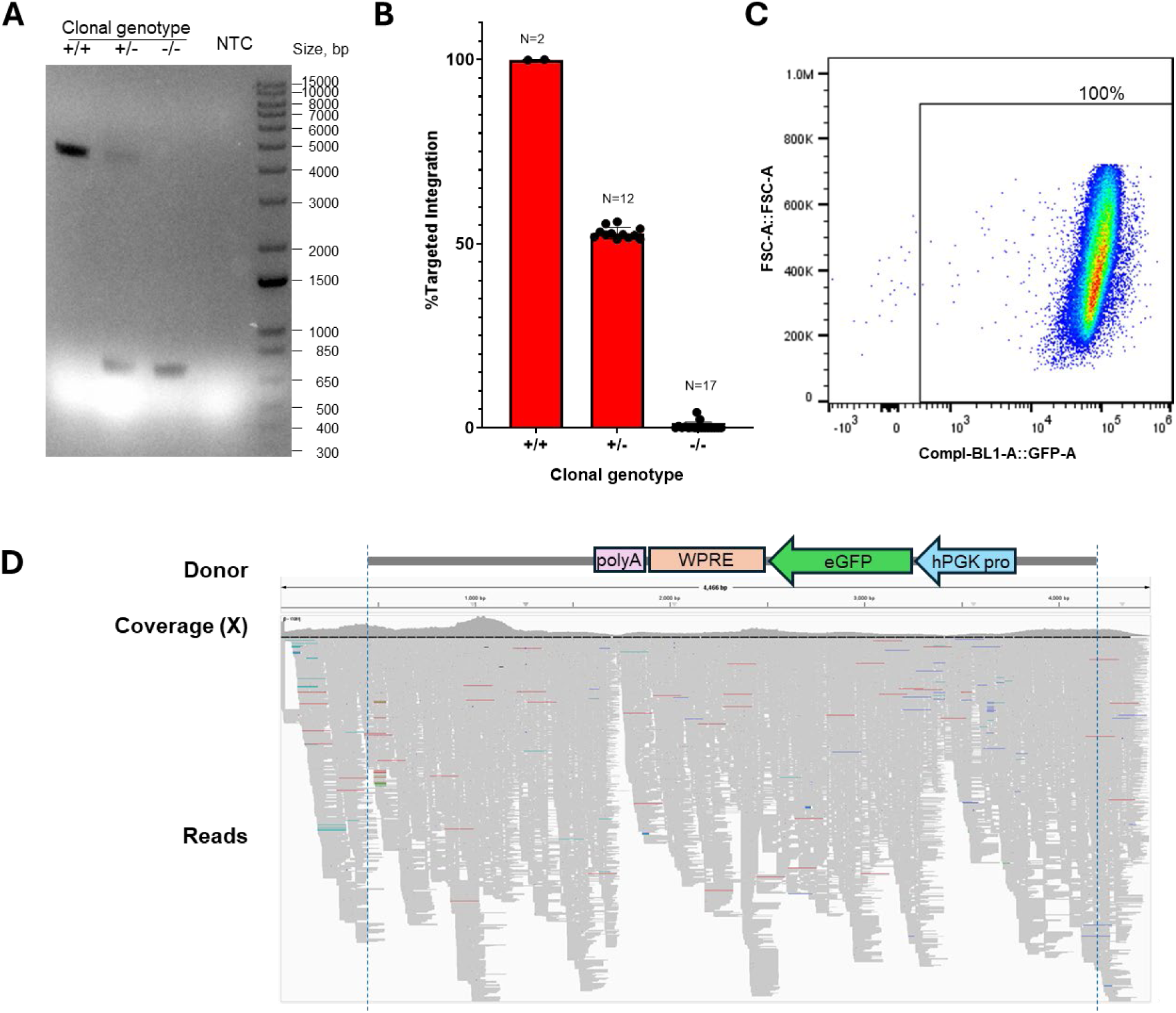
Targeted integration characterization in GFP positive single-cell clones. **a.** Representative results of PCR analysis of different types of single-cell clones. Note, the expected PCR product with full length TI, including a single copy donor and adjunct genomic sequence, is 4,465 bp, and the wild-type amplicon is 723 bp. NTC, no template control. P-ctrl, positive control. **b**. Results for all GFP-positive clones analyzed in this experiment. Each clone was characterized using our NGS TI assay and the %TI values are consistent with the results of the PCR assay. **c.** Flow cytometry of GFP expression in a biallelic clone. **d.** Nextera sequencing of PCR amplicon from the biallelic clone from panel c with minimum 500X coverage confirms the full-length integrated donor in the eeBxb1-ZFP biallelic clone.

**Extended Data Figure 11.**
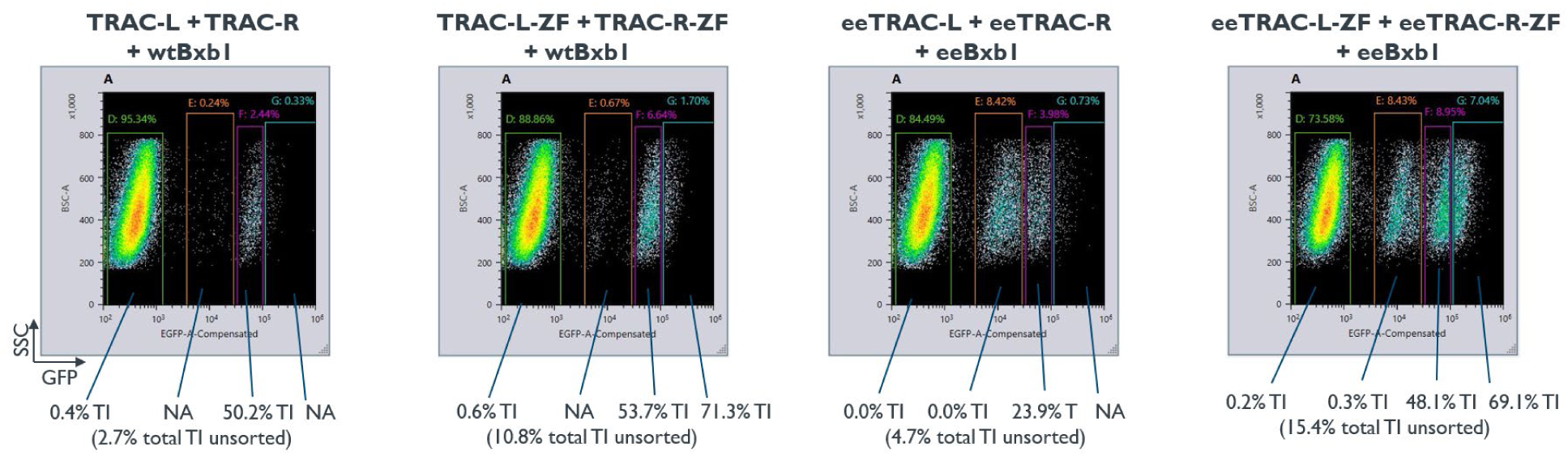
Targeted integration percentages in MINT-treated T cells sorted based on GFP expression levels. Nucleofected cells were sorted based on GFP expression into the 4 fractions indicated on the flow cytometry plots and TI percentages in the sorted fractions were determined. The percentage of cells in the different GFP expressing subpopulations is listed in the plots, the TI numbers are listed below the flow cytometry plots, with the number determined for the unsorted sample listed at the bottom in parentheses. NA indicates samples with too low of a cell number to yield a reliable result.

**Supplementary Figure 1.**
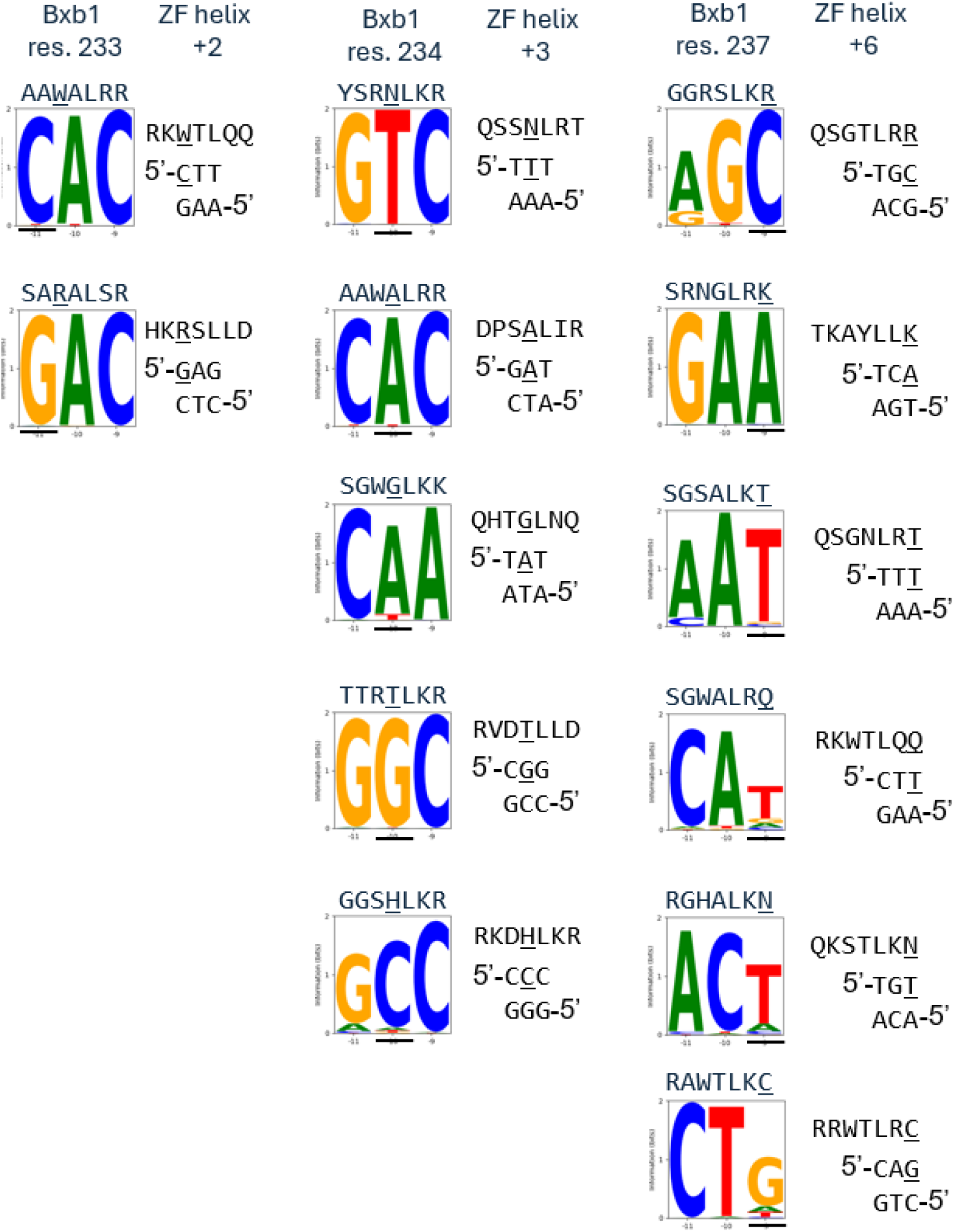
Similar Bxb1 helix and ZF helix interaction patterns with DNA. Bxb1 data from Figure 1f and ZF data from Ichikawa *et al.* (2023) except for QHTGLNQ which is from Wolfe *et al.* (2001). ZF data shown with primary DNA contacting strand on bottom to match Bxb1 data (QSGTLRR would bind 5’-GCA is standard ZF notation). Similar residue-base correlations for the Bxb1 helix and the ZF helix are underlined. Bxb1 helix and ZF helix patterns usually do not match at Bxb1 residue 231 and ZF helix position −1.

**Supplementary Figure 2.**
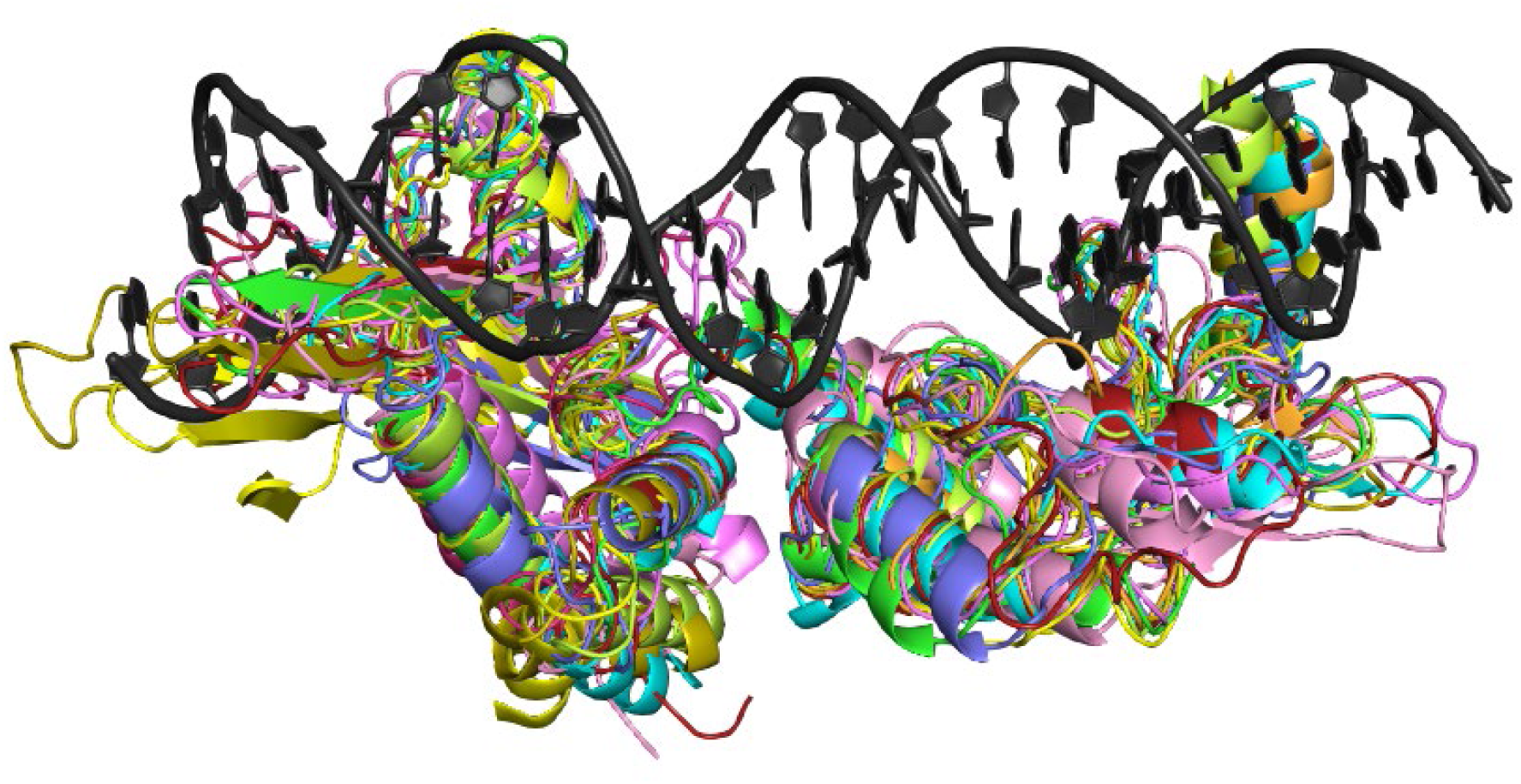
Structural alignment of predicted structures for RD and ZD domains of various LSIs. Shown are Pa01, Kp03, Nm60, Si74, BcyInt, and SscInt with the predicted structure of Bxb1. The structures of the predicted domains align well enough to identify the loop, helix, and hairpin regions of these LSIs as well as any LSIs that have a similar or better structural alignment with the predicted Bxb1 structure. Directed evolution using a residue randomization scheme at the same residues of the loop, helix, and hairpin of any such LSR should enable the identification of LSR variants that can target desired DNA sequences.

**Supplementary Figure 3.**
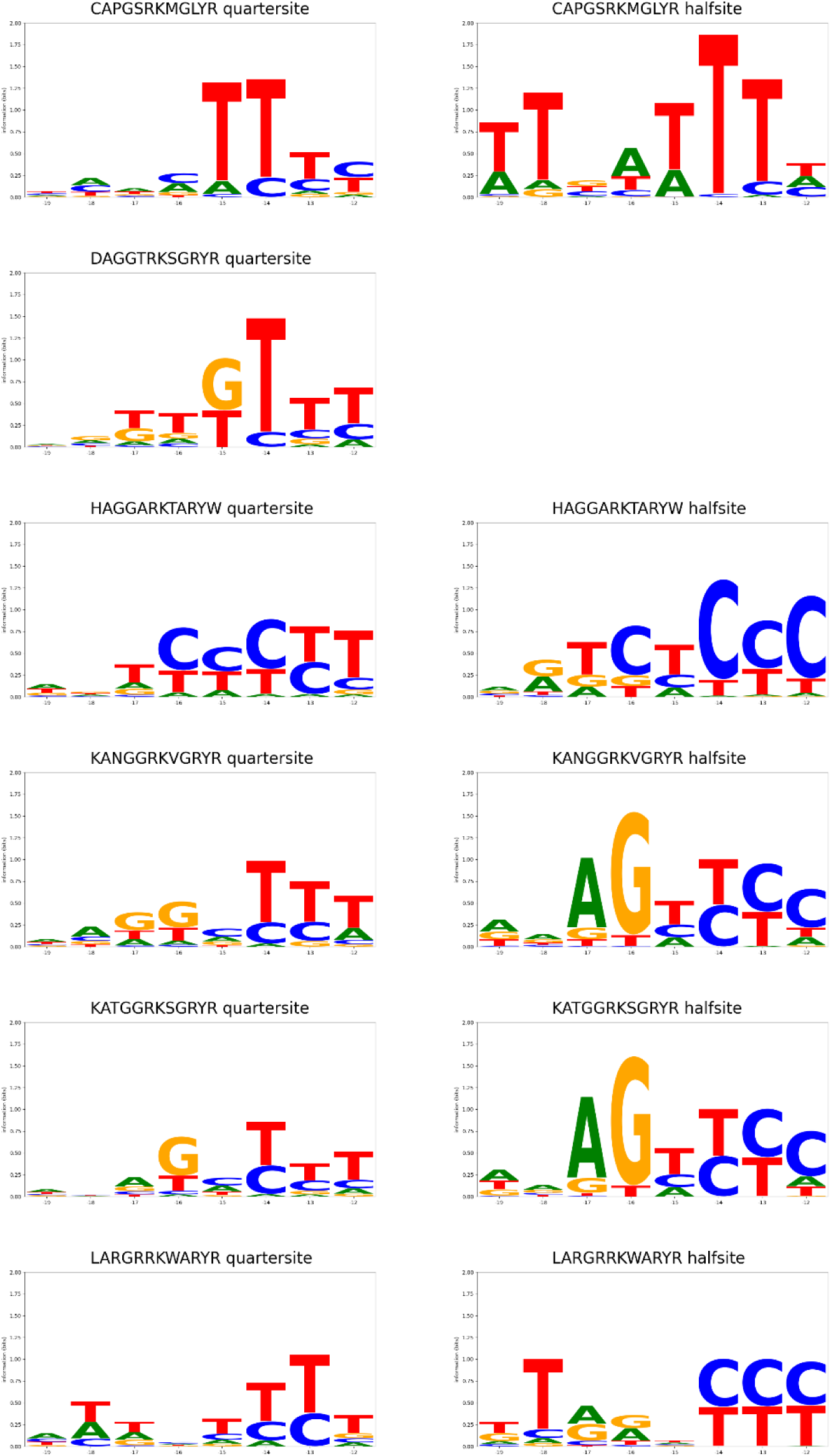
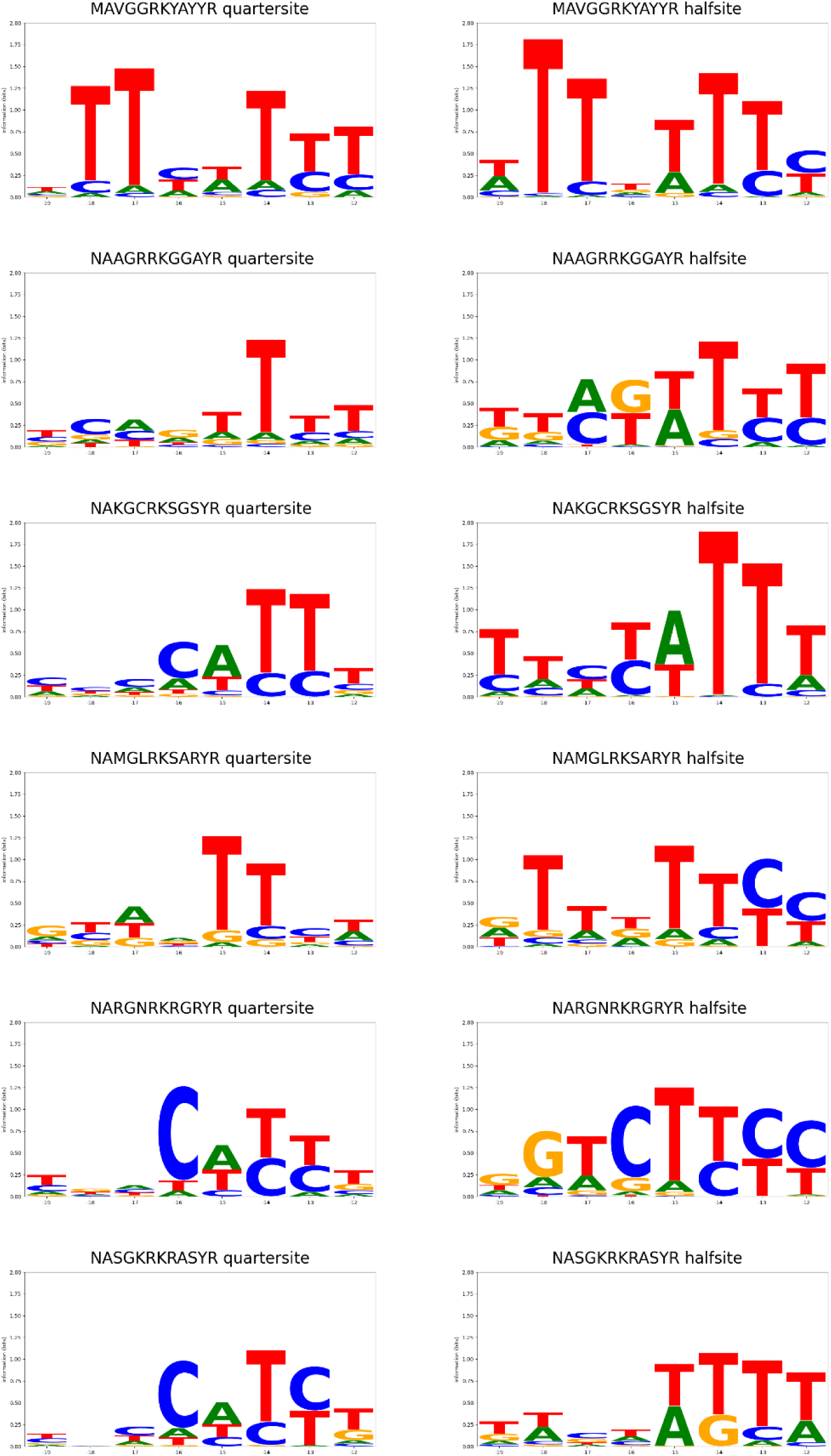
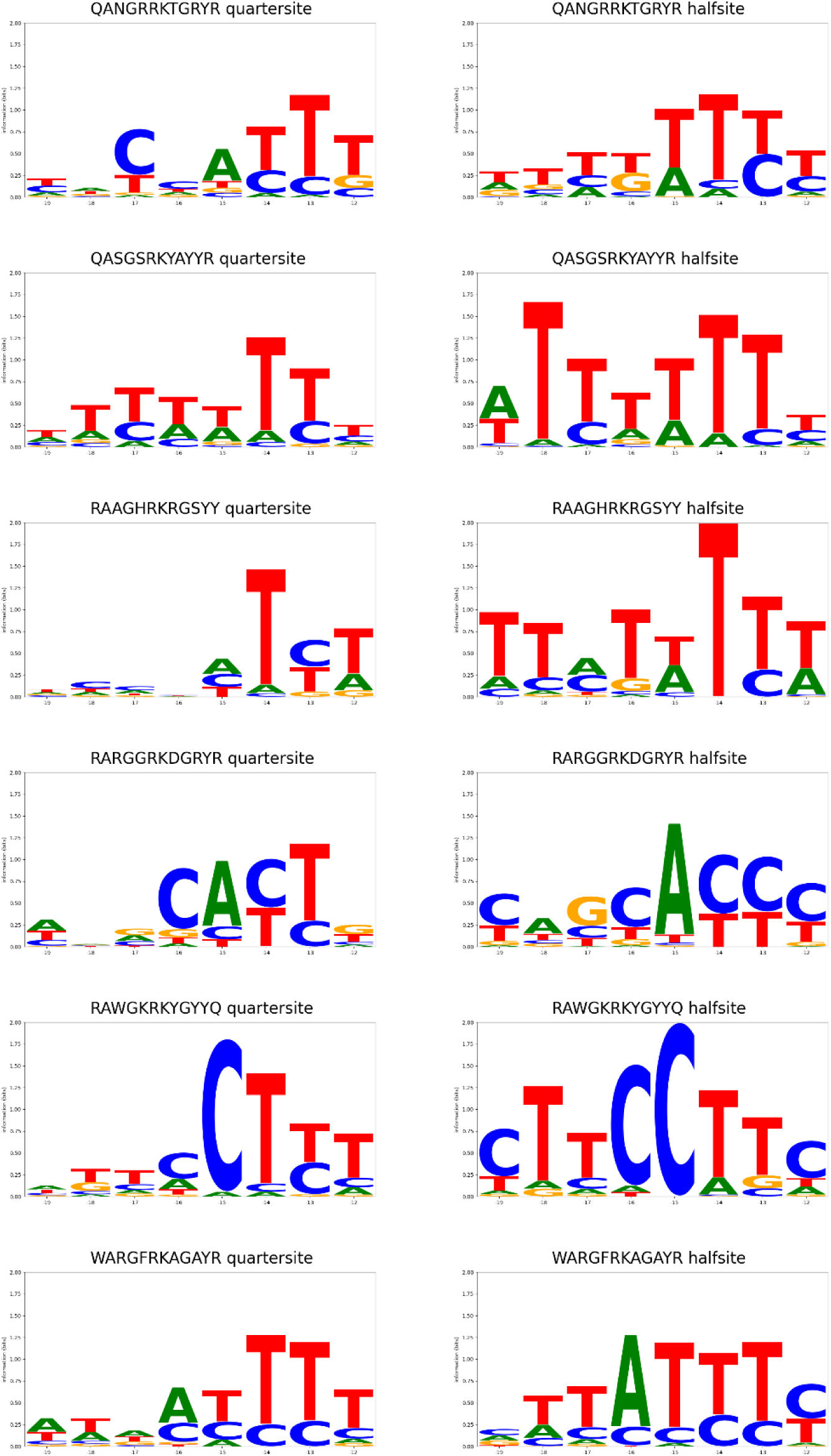
DNA target site preferences of engineered Bxb1 hairpins. DNA target site preferences for 18 engineered Bxb1 hairpins as measured in human K562 cells for recombination of a donor plasmid with two different libraries of DNA target site plasmids are plotted. Note that binding site preference data was not obtained for 8 hairpins in the hairpin archive table because these hairpins were only tested in combination with helix variants to target the endogenous sites indicated in the table. The “quarter-site” and “half-site” libraries are derived from 500 potential human targets for Bxb1 variants within disease related genes. The hairpin quarter-site pool has bases at positions −19 to −12 in the left half-site of attB varied to match the corresponding portion of the relevant target half-site (the right half-site of attB bears corresponding mutations at +12 to +19). Wild-type Bxb1 was included in the experiment to enable use of a single donor plasmid bearing a wild-type Bxb1 attP target so only sequences with read fractions at least 3 standard deviations above the read fraction for the wild-type Bxb1-only control at the same target sequence were analyzed. The sequence motif of the weighted average of the top 20 DNA targets at least 3 standard deviations above the wild-type-only control for positions −19 to −12 are plotted. If fewer than 20 sequences are above this cut-off then all successful sequences are plotted unless a single half-site represents more than half of the weighted average.

**Supplementary Figure 4.**
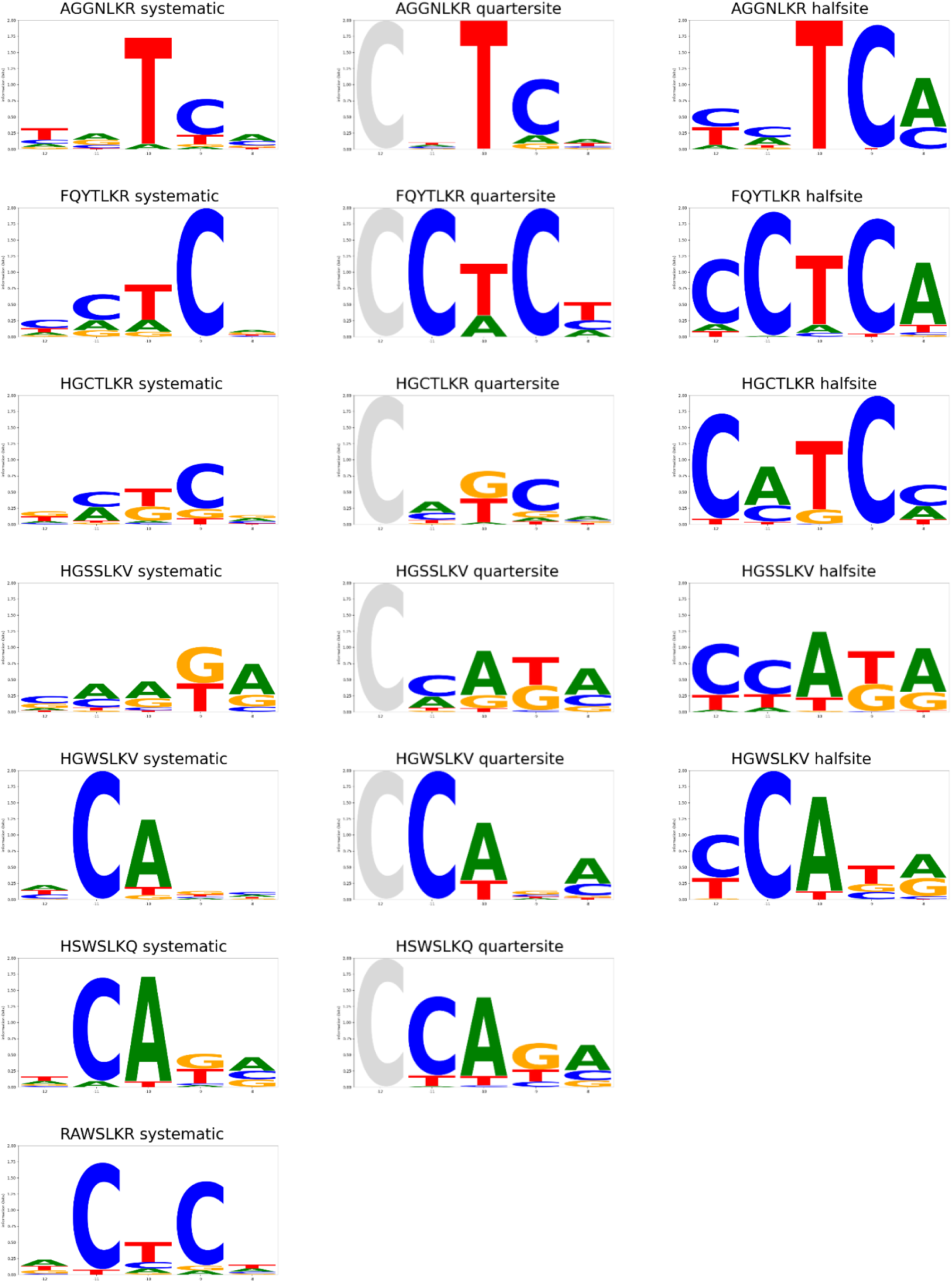
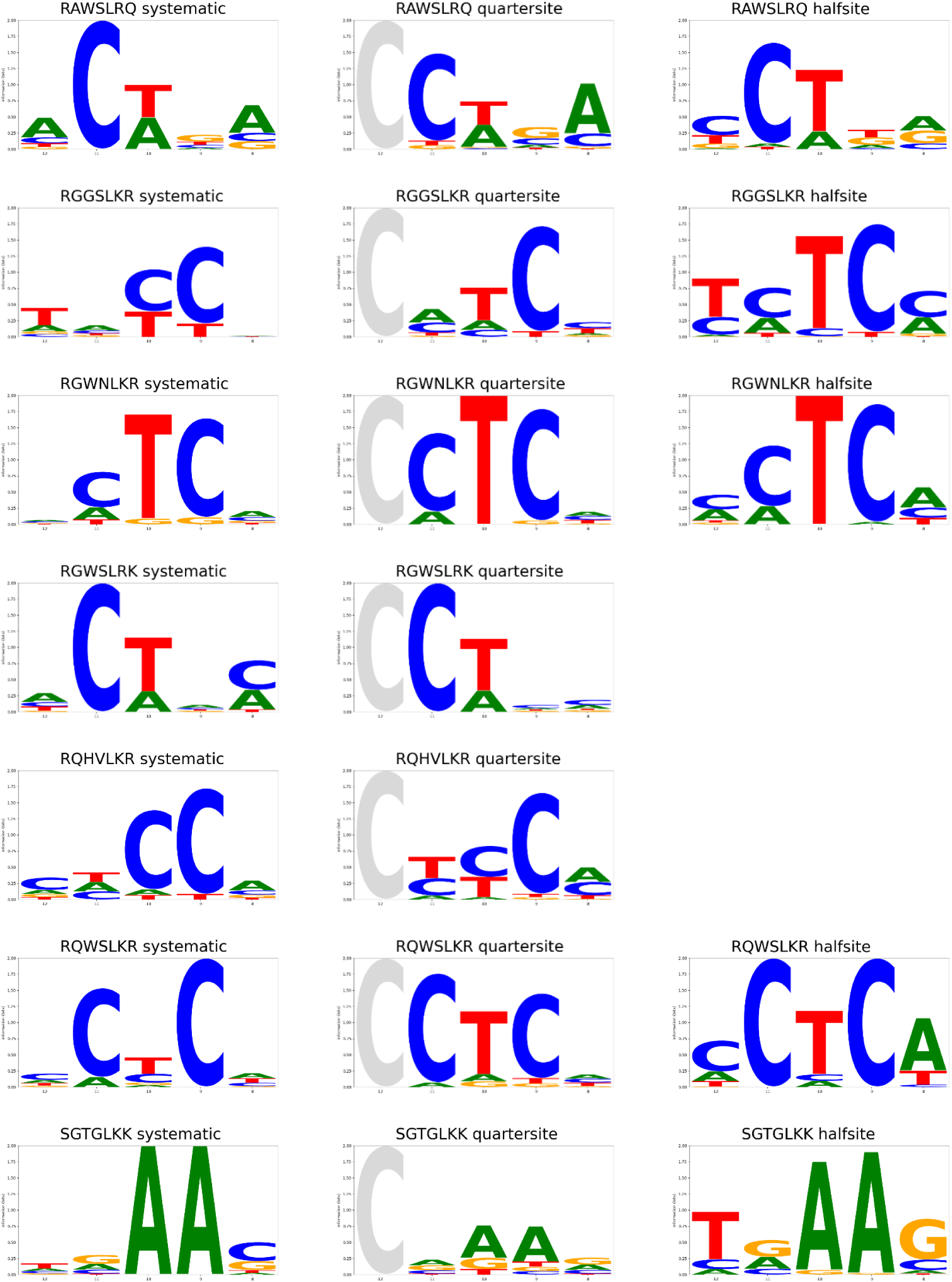
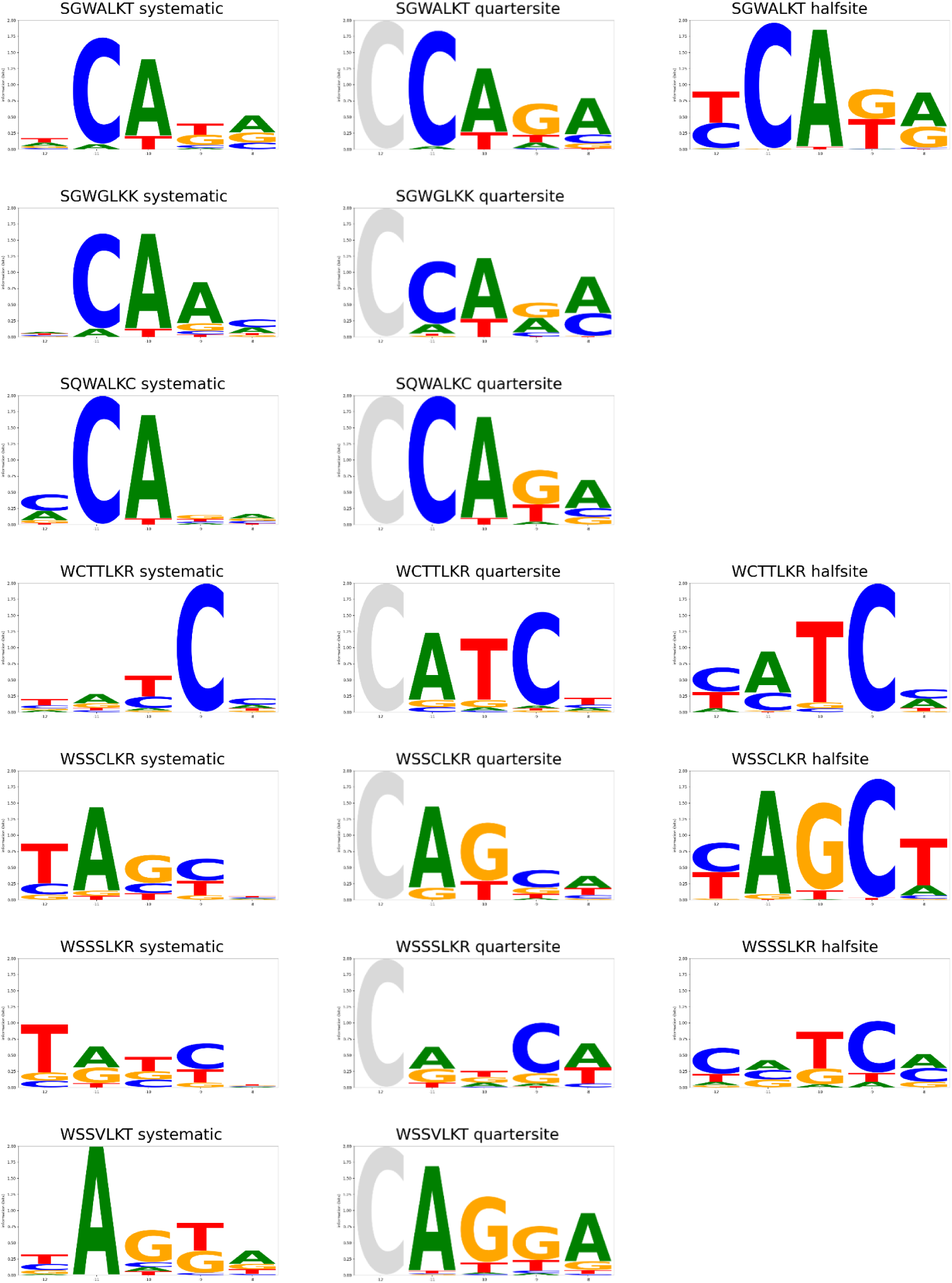
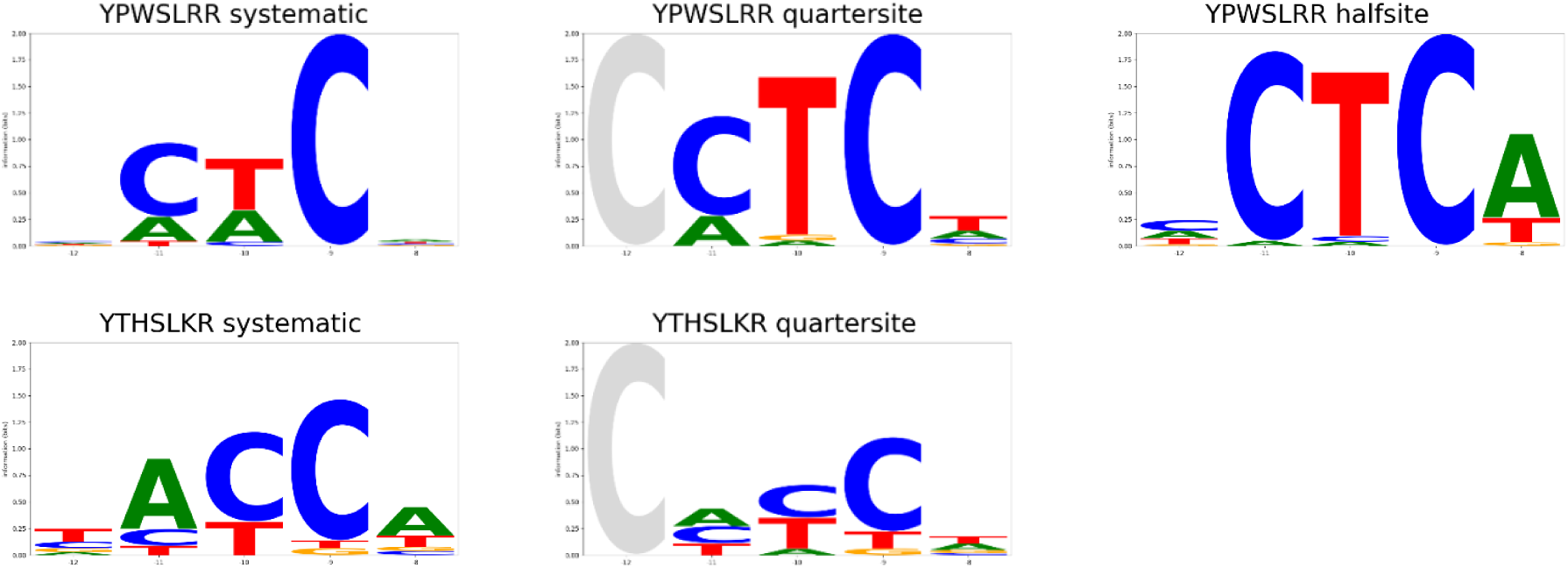
DNA target site preferences of engineered Bxb1 helices. DNA target site preferences for 23 engineered Bxb1 helices as measured in human K562 cells for recombination of a donor plasmid with three different libraries of DNA target site plasmids are plotted. The “systematic” target site library has bases at positions −12, −11, −10, −9, and −8 in the left half-site of attB fully randomized (the right half-site of attB bore corresponding mutations at +8, +9, +10, +11, and +12). Note that YGSALKQ was not characterized with these DNA target pools because it was only tested in combination with hairpin variants to target the endogenous TRAC locus. The “quarter-site” and “half-site” libraries are derived from 500 potential human targets for Bxb1 variants within disease related genes. Each member of the helix quarter-site library contains the sequence from −11 to −2 from one half-site of the 500 potential target sites (1000 library members in total) and an inverted repeat of this sequence at positions +2 to +11. The gray C indicates that the C at position −12 wasn’t varied in this library. The half-site libraries contain entire half-sites (−19 to −2) from these 500 potential target sites (1000 half-sites in total) with an inverted repeat of the same half-site from +2 to +19. All target sites in the library have the GT central dinucleotide (positions −1 and +1) from the wild-type attB target site in order to successfully recombine with the wild-type Bxb1 attP site in the donor. Wild-type Bxb1 was included in the experiment to enable use of a single donor plasmid bearing a wild-type Bxb1 attP target so only sequences with read fractions at least 3 standard deviations above the read fraction for the wild-type Bxb1-only control at the same target sequence were analyzed. The sequence motif of the weighted average of the top 20 DNA targets at least 3 standard deviations above the wild-type only control for positions −12, −11, −10, −9, and −8 are plotted. If fewer than 20 sequences are above this cut-off then all successful sequences are plotted unless a single half-site represents more than half of the weighted average.

